# Investigating molecular mechanisms of 2A-stimulated ribosomal pausing and frameshifting in *Theilovirus*

**DOI:** 10.1101/2020.08.11.245068

**Authors:** Chris H. Hill, Georgia M. Cook, Sawsan Napthine, Anuja Kibe, Katherine Brown, Neva Caliskan, Andrew E. Firth, Stephen C. Graham, Ian Brierley

## Abstract

The 2A protein of Theiler’s murine encephalomyelitis virus (TMEV) acts as a switch to stimulate programmed −1 ribosomal frameshifting (PRF) during infection. Here we present the X-ray crystal structure of TMEV 2A and define how it recognises the stimulatory RNA element. We demonstrate a critical role for bases upstream of the originally predicted stem-loop, providing evidence for a pseudoknot-like conformation and suggesting that the recognition of this pseudoknot by beta-shell proteins is a conserved feature in cardioviruses. Through examination of PRF in TMEV-infected cells by ribosome profiling, we identify a series of ribosomal pauses around the site of PRF induced by the 2A-pseudoknot complex. Careful normalisation of ribosomal profiling data with a 2A knockout virus facilitated the identification, through disome analysis, of ribosome stacking at the TMEV frameshifting signal. These experiments provide unparalleled detail of the molecular mechanisms underpinning *Theilovirus* protein-stimulated frameshifting.

## Introduction

Cardioviruses are a diverse group of picornaviruses that cause encephalitis, myocarditis and enteric disease in a variety of mammalian hosts including rodents, swine and humans (1). The *Cardiovirus B* or *Theilovirus* species comprises several isolates including Sikhote-Alin virus (2), rat theilovirus and Theiler’s murine encephalomyelitis virus (TMEV), all of which are genetically distinct from the *Cardiovirus A* species encephalomyocarditis virus (EMCV) (3). Within the *Cardiovirus B* species, TMEV has been extensively characterised and serves as a mouse model for virus-induced demyelination and multiple sclerosis (4). Like all picornaviruses, TMEV replication is cytoplasmic and begins with the translation of its single-stranded ∼8 kb positive-sense RNA genome. The resultant polyprotein (L-1ABCD-2ABC-3ABCD) is subsequently processed, mainly by the virally encoded 3C protease (5, 6).

Several “non-canonical” translation events occur during the production of the TMEV polyprotein. First, initiation is directed by a type II internal ribosome entry site (IRES) in the 5′ untranslated region (UTR) (7, 8). Secondly, a co-translational StopGo or ribosome “skipping” event occurs at the junction between the 2A and 2B gene products (9, 10). In this process, the peptidyl-transferase reaction fails between the glycine and proline in a conserved D(V/I)ExNPG|P motif, releasing the upstream L-1ABCD-2A product as the ribosome continues translating the downstream 2BC-3ABCD region. Note that in TMEV, however, the presence of a 3C protease cleavage site near the start of the 2B protein appears to make the StopGo reaction functionally redundant (11, 12). Thirdly, programmed −1 ribosomal frameshifting (PRF) diverts a proportion of ribosomes out of the polyprotein reading frame and into a short overlapping ORF, termed 2B*, near the start of the 2B protein. In TMEV this ORF is only eight codons in length and the resulting transframe product 2B* has just 14 amino acids (∼ 1.4 kDa), with no established functional role (11). Thus it has been hypothesised that in TMEV, the main function of PRF may simply be to downregulate translation of the enzymatic proteins encoded downstream of the frameshift site, particularly in the late stages of infection (11).

PRF is common amongst RNA viruses, where it is used as a translational control strategy to express gene products in optimal ratios for efficient virus replication. Additionally, the utilisation of overlapping ORFs permits more information to be encoded by a small genome. Our mechanistic understanding of PRF has been informed by the study of examples in hundreds of viruses (reviewed in ref. 13-15). Generally, PRF involves two elements within the viral messenger RNA (mRNA). A heptameric shift site or “slippery sequence” of the form X_XXY_YYZ (where XXX represents any three identical nucleotides or certain other triplets such as GGU, YYY represents AAA or UUU, and Z is any nucleotide except G (16) is located 5 – 9 nucleotides upstream of a structured RNA “stimulatory element” (usually a stem-loop or pseudoknot) that impedes the progress of the elongating ribosome, such that the ribosome pauses with the shift site in the decoding centre (17–19). This can facilitate a change of reading-frame if the codon-anticodon interactions of the P-site and A-site tRNAs slip and recouple in the −1 frame (XXY → XXX and YYZ → YYY, respectively) during resolution of the stimulatory element. For any given system, such PRF generates a fixed ratio of products set by parameters that include the energetics of codon-anticodon pairing at the shift site, the conformational flexibility of the stimulatory element and the resistance of this element to unwinding by the ribosomal helicase (20, 21). In cardioviruses, however, PRF exhibits some intriguing mechanistic exceptions. First, the conserved G_GUU_UUU shift site is located 13 – 14 nucleotides upstream of the stimulatory stem-loop, seemingly too far away to position the shift site in the decoding centre during a pause. Secondly, the viral 2A protein is required as a *trans*-activator of frameshifting in cells and *in vitro* (11,22–24) which permits temporally controlled, variable-efficiency frameshifting related to the amount of 2A protein in the cell during infection. To date, cardioviruses present one of only two known examples of protein-stimulated frameshifting, along with porcine reproductive and respiratory syndrome virus (PRRSV) and other arteriviruses (family *Arteriviridae*), where a complex of viral nsp1β and host poly(C) binding protein stimulates PRF (25–27).

Our previous investigations of protein-stimulated frameshifting in EMCV and TMEV have revealed that the viral 2A protein acts in complex with a stem-loop downstream of the slippery sequence, and that this interaction can be disabled by mutating either a conserved cytosine triplet in the loop or a pair of conserved arginine residues in the protein (11,22,23). More recently, we solved the structure of EMCV 2A, revealing a novel protein fold that permits binding to both the stem-loop and to ribosomal RNA with high affinity (28). However, 2A protein sequences are highly divergent within cardioviruses, and the TMEV protein shares only ∼ 27% pairwise amino acid sequence identity with its EMCV orthologue. Additionally, the stem-loop that comprises the stimulatory element in TMEV is significantly more compact than the equivalent structure in EMCV. Here we present the X-ray crystal structure of TMEV 2A and investigate the interaction with its cognate RNA using a variety of biochemical and biophysical techniques. We define a minimal TMEV stimulatory element necessary for 2A binding and show that the protein forms a 1:1 RNA-protein complex with nanomolar affinity. We provide evidence that the alternative pseudoknot-like conformation recently described for the EMCV stimulatory element (28) is also likely to be present in TMEV and other cardioviruses. Finally, we use metabolic labelling and ribosome profiling to study 2A-stimulated frameshifting and translation of the TMEV genome at sub-codon resolution in infected cells. Together, this body of work defines the molecular basis of *Theilovirus* protein-stimulated frameshifting, one of the most efficient frameshifting events known in nature.

## Results and Discussion

### Structure of TMEV 2A reveals a beta-shell protein with a divergent RNA-binding surface

TMEV 2A protein was purified following recombinant expression as a GST fusion in *E. coli* (Figure 1A). After cleavage of the GST tag and removal of contaminating nucleic acids by heparin affinity chromatography, high-salt conditions were required to prevent aggregation and maintain solubility. Under optimised conditions, size-exclusion chromatography coupled to multi-angle light scattering (SEC-MALS) revealed the protein to be predominantly monomeric (Figure 1B, peak 2) with an observed M_w_ of 16,683 Da compared to a theoretical mass of 15,941 Da, as calculated from the primary sequence. A small proportion of trimers was also present (Figure 1B, peak 1: observed mass of 46,867 Da vs. theoretical mass of 47,823 Da). We crystallised the protein but were unable to solve the structure by molecular replacement using the crystal structure of the closest homologue, EMCV 2A (28), as a search model. Instead, we obtained experimental phases via bromine derivatisation and determined the structure by single-wavelength anomalous dispersion analysis. The asymmetric unit of the cubic cell contained a single copy of 2A, which was refined to 1.9 Å resolution (Table S1, Figure 1C). Interpretable electron density was visible for 2A residues 1-126, with a short G_-4_P_-3_L_-2_G_-1_S_0_ N-terminal extension that resulted from 3C protease cleavage of the GST tag. In the crystalline lattice these residues mediate contacts between symmetry-related molecules, consistent with recombinant 2A forming trimers at high concentrations (Figure S1). These residues project away from the globular TMEV domain and lack regular secondary structure, suggesting that they will be flexible at lower concentrations of 2A in solution where monomers predominate (Figure 1B). Since tag-derived residues mediate the inter-subunit contacts, this 2A trimerisation is unlikely to be physiologically relevant.

**Figure 1.**
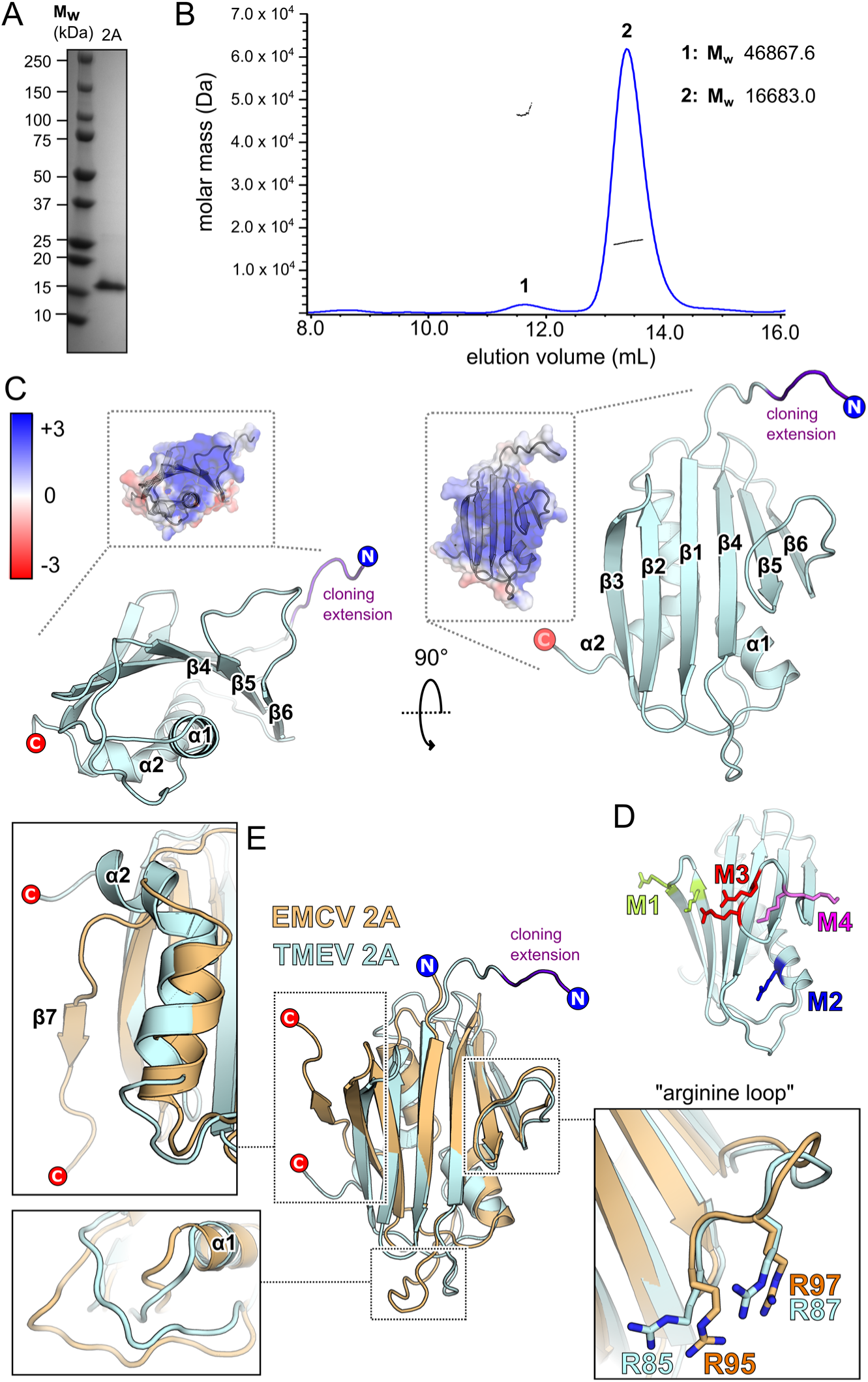
TMEV 2A adopts the beta-shell fold. **A.** SDS-PAGE analysis of TMEV 2A after Ni-NTA, heparin affinity and size-exclusion chromatography. The gel was stained with Coomassie blue. **B.** SEC-MALS analysis of 2A in high-salt buffer at 3.1 mg/mL. The differential refractive index curve (blue) is shown and the weight-averaged molar masses for indicated peaks are listed. **C.** X-ray crystal structure of TMEV 2A in two orthogonal views. N- and C-termini are indicated, and amino acids introduced as a result of the cloning strategy are labelled in purple. *<INSET>* Electrostatic surface potential calculated at pH 7.4, coloured between +3 (blue) and −3 kT/e^-^ (red). **D.** Locations of mutations are colour-coded and shown as sticks: M1 (K24A / R28A, green); M2 (R45A, blue); M3 (R85A / R95A, red); M4 (K90A / K91A, pink). **E.** Superposition of TMEV 2A (blue) with EMCV 2A (orange). *<INSETS<* Close-up views of regions of interest including the divergent structure at the C-terminus, the longer α1-β4 loop present in EMCV 2A and the conserved arginine loop.

The structure of TMEV 2A reveals a globular β_3_αβ_3_α fold, with a similar “beta-shell” architecture to EMCV 2A (C_α_ backbone RMSD of 2.65 Å over 130 residues) (28). An extensive curved antiparallel six-stranded beta sheet packs against two alpha helices on the concave surface of the sheet, whilst the loops between adjacent beta strands project from the opposite convex face. The protein is highly basic (pI ∼ 9.4) and most of the contributing lysine, arginine and histidine residues are solvent-exposed. At physiological pH the protein will thus have a positive electrostatic surface potential across the convex face of the beta sheet, surrounding loops and the N-terminal end of helix α1, suggesting a putative RNA-binding surface (Figure 1C). This is supported by a previous biochemical analysis of TMEV 2A function, in which we made several point mutations of conserved basic residues and assessed their ability to stimulate PRF (23) (Figure 1D). The M3 mutant (R85A / R87A, termed ‘2A-mut’ in Napthine *et al.*, 2019) was found to completely inhibit frameshifting, and the M1 mutant (K24A / R28A) was found to reduce it by approximately four-fold. Both M3 and M1 are in surface loops on either side of this large, positively charged beta sheet, consistent with a role for this face of the protein in forming electrostatic interactions with the ribose phosphate backbone of the PRF stimulatory element. In contrast, M2 (R45A) had no effect – indeed most of this residue is buried (only approximately 11.6% of the residue’s surface area being solvent-accessible) and it likely plays a structural role in stabilising packing of helix α1 against the underside of the central sheet. Perhaps surprisingly, M4 (K90A / K91A) also had no effect, despite these residues being located in the same positively charged loop as the essential R85 and R87. This demonstrates that precise spatial positioning of charge within this loop is key to the specificity of RNA-binding.

Despite low sequence identity, the overall fold of TMEV 2A is similar to EMCV 2A (Figure 1E). The most functionally important part of EMCV 2A is the “arginine loop” between strands β5 and β6 – necessary for PRF (22), RNA binding (28) and nuclear localisation (29). The two arginines in this loop (R95 / R97 in EMCV 2A; R85 / R87 in TMEV 2A) are amongst the only surface-exposed residues that are completely conserved across both species of cardiovirus. (Figure S2A and B). Structurally, this loop adopts an almost identical conformation, consistent with mutagenesis data suggesting that they are functionally equivalent (22, 23). However, there are also some key differences between the two 2A orthologues (Figure 1E). The loop region between the end of α1 and the start of β4 is much longer in EMCV 2A. This loop, and the N-terminal end of α1, are two of three regions that contribute to the RNA-binding surface in EMCV 2A (28). These two regions are not conserved in TMEV 2A (Figure S2C): the backbone geometry is different, and, with the exception of K63 and R45, there are no chemically equivalent side chains in the vicinity that could form similar interactions. To investigate this further, we prepared variants of TMEV 2A containing point mutations (R85A / R87A, K63A, K83A and K53A / H56A; Figure S3A) and tested their ability to bind stimulatory element RNA (Figure S3B). Only the R85A / R87A protein showed a significant defect compared to wild-type protein, confirming the unique importance of this conserved arginine loop. Beyond this, the RNA binding surface in TMEV 2A likely differs from EMCV 2A, perhaps involving additional residues on the surface of the beta sheet that are only conserved amongst *Theilovirus* isolates (e.g. R7, D9, K24; Figure S2A-C). This is supported by a reduction in PRF seen with the M1 (K24A) mutant (23).

The conservation at the C-terminus of the protein (Figure S2A and B) is concentrated in the D(V/I)ExNPG motif required for StopGo peptide release between 2A and 2B gene products in other cardioviruses (9, 10). As expected, this motif is unstructured in both proteins, and if we consider only the ordered amino acids, we observe some of the largest structural differences (Figure 1E). In EMCV 2A, the shorter α2 helix leads to a more pronounced curvature of the central beta sheet, and the C-terminus forms a short β7 strand that packs against β3. Conversely, in TMEV 2A, the α2 helix continues with a pronounced 90° kink until the end of the protein. This has consequences for the structure of the putative YxxxxLΦ motif at the C-terminus that has previously been reported to sequester eIF4E in a manner analogous to 4E-BP, thereby disabling cap-dependent translation of host mRNA (29). In TMEV 2A, the first tyrosine residue of this motif (Y119) is located on the buried side of the α2 helix (with only approximately 4.0% of the residue’s surface area being solvent accessible) and is therefore not available to interact with eIF4E. A second tyrosine (Y120) is more exposed (approximately ∼ 26.7% of the residue’s surface area being solvent accessible), but given the local secondary structure, it is unlikely that L124 and I125 would be able to interact with eIF4E in the same way as 4E-BP without a significant conformational change (Figure S2D). In EMCV 2A, the putative YxxxxLΦ motif is not helical, instead forming a more extended backbone (28). Superposition of this region reveals that this motif is not structurally conserved between 2A orthologues (Figure S2E). Therefore, the interaction with eIF4E must either involve other parts of the protein or must be accompanied by significant structural rearrangement of the α2 helix of TMEV.

### A minimal 37 nt pseudoknot in the viral RNA is required for 2A binding

The PRF stimulatory element in TMEV consists of a stem-loop (seven base-pair stem and 10 nucleotide loop) located 14 nucleotides downstream from the shift site (Figure 2A). This is more compact than the equivalent element in EMCV, which has a 21 nt loop that may contain an additional stem (22, 28). However, in both viruses, three conserved cytosines in the loop are essential for 2A binding (22, 23). To determine the minimal RNA element necessary for interaction with TMEV 2A, we prepared a series of synthetic RNA constructs (Figure 2B) and assessed 2A binding by electrophoretic mobility shift assays (EMSA; Figure 2C) and microscale thermophoresis (MST; Figure 2D).

**Figure 2.**
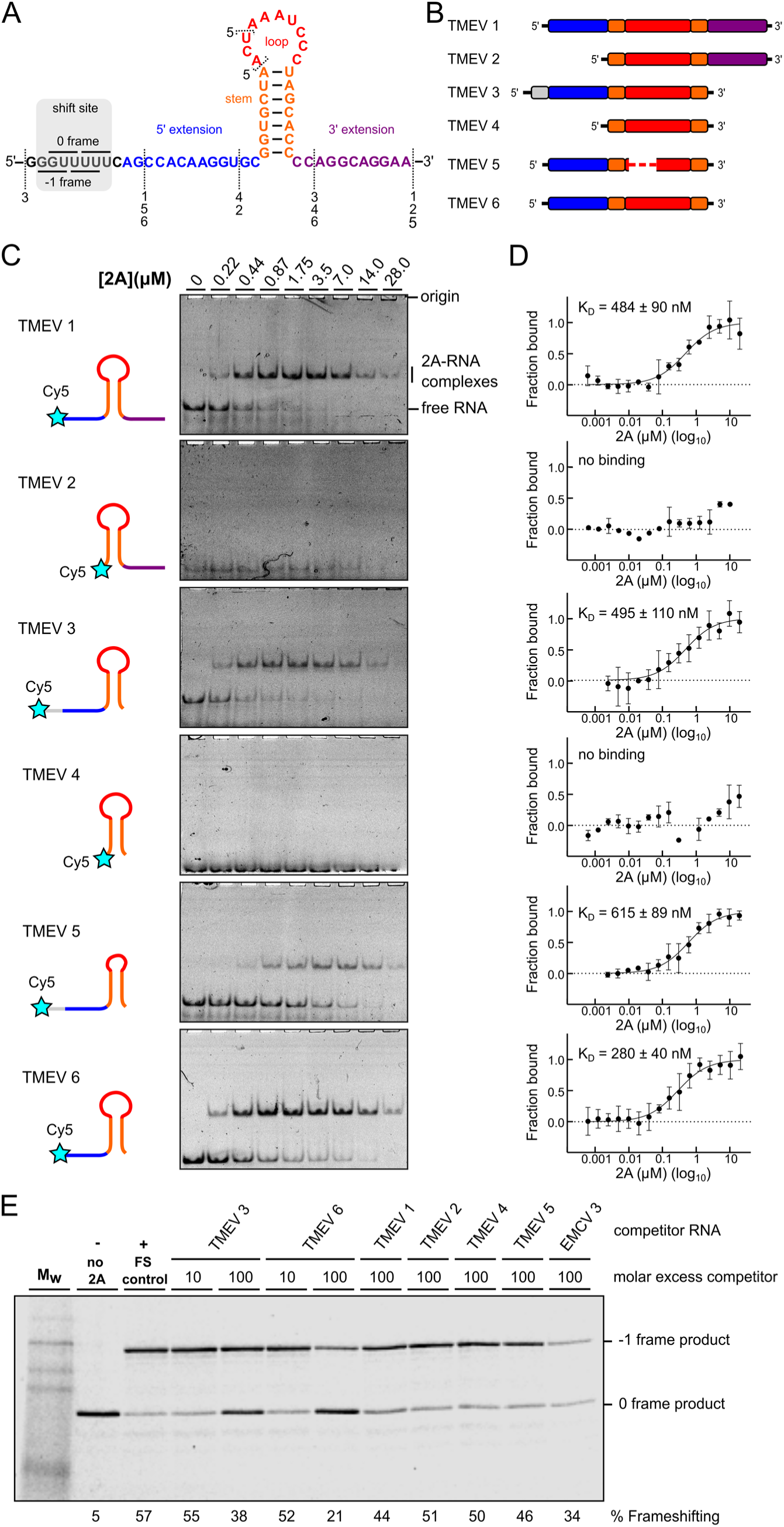
2A recognises a minimal 37 nt stimulatory element in the viral RNA. **A, B.** Sequences and schematic diagrams of the TMEV 1-6 constructs used to assay 2A binding. **C.** EMSA analysis conducted with 50 nM Cy5-labelled RNAs and 2A concentrations between 0 and 28 μM. Following non-denaturing electrophoresis, fluorescence was imaged using a Typhoon scanner. **D.** MST binding curves and apparent K_D_ values using the same constructs. **E.** Experiment showing the effects of titrating excess short RNAs (TMEV 1-6) as competitors into an *in vitro* frameshift reporter assay. The concentrations of the reporter mRNA and 2A were kept constant in the RRL and short RNAs were added in 10-and 100-fold molar excess relative to the reporter mRNA, as indicated. Translation products were visualised by using autoradiography, and % frameshifting was calculated following densitometry and correction for the number of methionines present in 0 frame and −1 frame products.

Binding of 2A was generally high affinity, with dissociation constants in the sub-micromolar range. (Figure 2D). Truncation of either the shift-site (TMEV 1, K_D_ = 484 ± 90 nM), the 3′ extension (TMEV 3, K_D_ = 495 ± 110 nM) or both (TMEV 6, K_D_ = 280 ± 40 nM) had little effect on 2A binding. Shortening the loop by three nucleotides (TMEV 5, K_D_ = 615 ± 89 nM) slightly reduced the affinity, however removal of the 5′ extension (TMEV 2, TMEV 4) completely abolished 2A binding, even though the stem-loop in TMEV 4 is predicted to be intact, and the three essential cytosines are present in the loop. To validate that these small RNAs are adopting conformations relevant to PRF, we performed competition experiments. A dual-luciferase-based reporter mRNA containing the TMEV shift site and stimulatory element was prepared and designed such that 0 frame and −1 frame products would be easily distinguishable by SDS-PAGE following *in vitro* translation in rabbit reticulocyte lysates (23). In the presence of 2A, PRF occurred efficiently (∼ 57%; Figure 2E), but inclusion of a molar excess of the small RNA would be predicted to reduce PRF efficiency if the competitor RNA were to sequester 2A from the reporter mRNA. In line with the EMSA assays, TMEV 3 and TMEV 6 both efficiently competed with the reporter mRNA, reducing PRF to 38% and 21%, respectively. RNAs lacking the 5′ extension (TMEV 2, TMEV 4) were unable to compete (Figure 2E).

We have recently shown that the 5′ extension is also important for 2A binding in EMCV, where it likely forms the second stem of an RNA pseudoknot via interaction with the CCC loop motif (28). However, given that the equivalent loop sequence is 11 nt shorter in TMEV, it was unclear whether this alternate conformation would be topologically possible for the more compact RNA element. Nevertheless, an alignment of cardiovirus RNA sequences that direct PRF shows that, in addition to the cytosine triplet, there are several nucleotides in the 5′ extension that are completely conserved in all isolates (Figure 3A, Figure S4). To investigate this in more detail, we truncated the 5′ extension one nucleotide at a time (Figure 3B) and assessed 2A binding by EMSA (Figure 3C). Removal of the first six nucleotides of the 5′ extension (CAGCCA; TMEV 7–9) had no dramatic effect on 2A binding, however removal of nucleotides from the conserved CAAGG motif progressively reduced binding until it was undetectable (TMEV 12). To explore this further, we made point mutations and assessed their effects in a frameshift reporter assay (Figure 3D). As expected, loop mutations C36G or C38G abolished PRF. However, frameshifting in the C38G background was restored by introducing a G17C mutation in the 5′ extension, demonstrating the necessity for a base pair between positions 17 and 38. The importance of this alternate conformation for 2A recognition was verified by EMSA analysis (Figure 3E). In RNA TMEV 9, which comprises the minimal functional element as defined by the deletion analysis, individual mutation of either C38 or G17 completely abrogated 2A binding. An unrestrained RNA folding simulation (30) identified a pseudoknot-like conformation consistent with the biochemical data, including a base pair between G17 and C38 (Figure 3F). This is consistent with our evidence for a pseudoknot-like conformation in the EMCV sequence that is selectively recognised by 2A (28). Whilst the double C38G+G17C mutant could rescue PRF in our *in vitro* reporter assay, it did not restore 2A binding by EMSA, either in the background of TMEV 9 (Figure 3E) or the longer TMEV 1 RNA (data not shown). It is possible that the double mutant forms a topologically equivalent conformer that is either less stable and/or binds 2A with lower affinity. Thus, whilst this RNA is still able to stimulate PRF, the association does not persist for the necessary timescales to permit observation in a dissociative technique such as EMSA.

**Figure 3.**
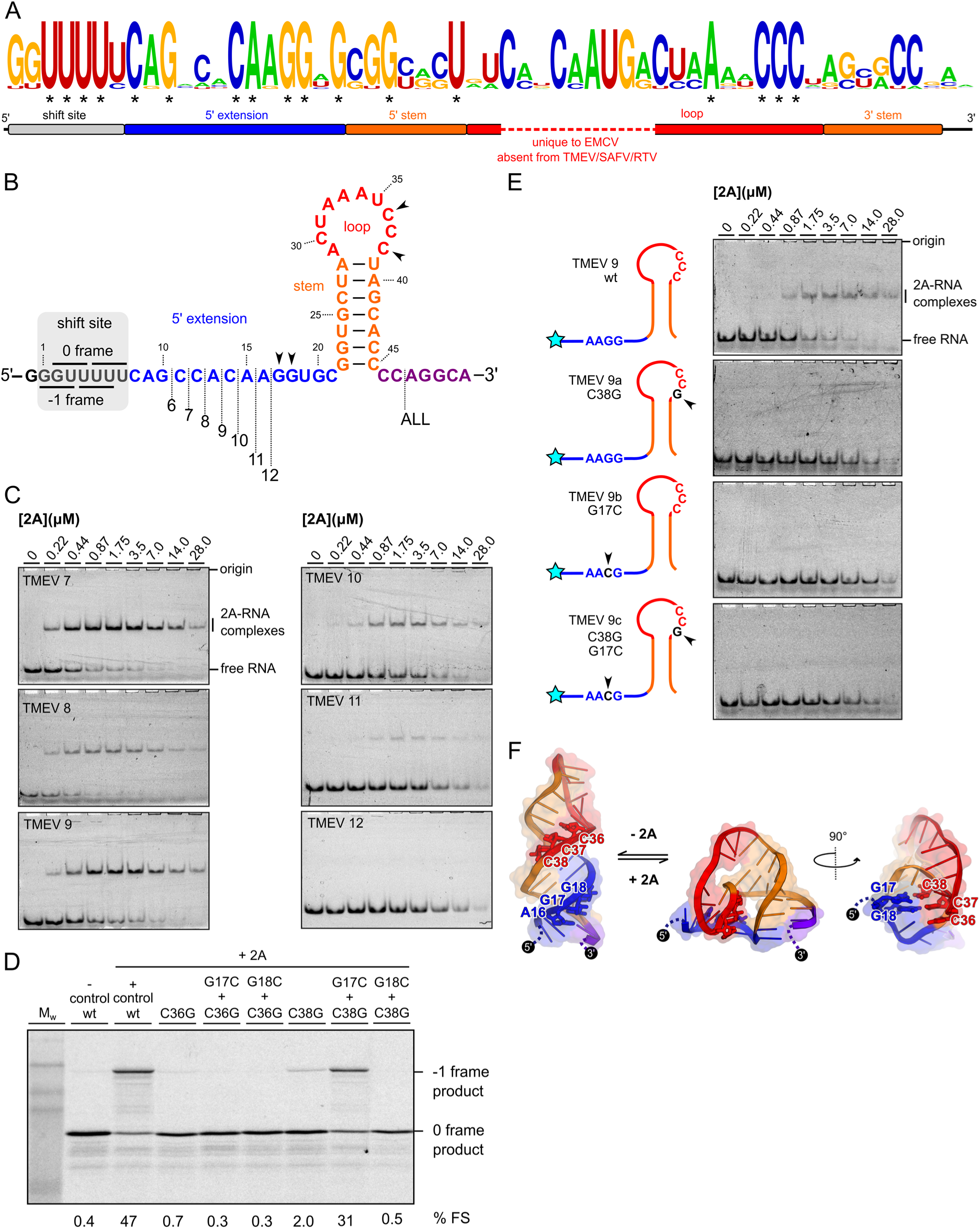
A conserved AAGG motif present in the 5′ extension is required for 2A binding. **A.** Logo plot showing conservation of aligned cardiovirus RNA sequences (see Figure S4) spanning the experimentally defined minimal element. Asterisks indicate invariant nucleotides. **B.** Schematic diagrams of the TMEV 7-12 sequences used to assay 2A binding. **C.** EMSA analysis conducted with 50 nM Cy5-labelled RNAs and 2A concentrations between zero and 28 μM. Following non-denaturing electrophoresis, fluorescence was imaged using a Typhoon scanner. **D.** Frameshift reporter assays showing that mutation of the loop CCC motif inhibits frameshifting, but complementary mutations in the 5′ AAGG motif that allow base pairing between positions 17 and 38 restore frameshifting. **E.** EMSA analysis showing attenuation of 2A binding to the minimal TMEV 9 RNA by introducing point mutations to either the 5′ AAGG motif or loop CCC motif. **F.** Hypothetical equilibrium between predicted stem-loop and alternate pseudoknot-like conformations, colour-coded as in A. The pseudoknot-like conformation involves a base-pair between G17 and G18 in the 5′ extension, and C38 and C37 in the loop (shown as sticks).

To assess the binding affinity and stoichiometry in solution with unlabelled RNA, we carried out isothermal titration calorimetry (ITC) experiments with TMEV 6 and 2A protein. To prevent protein aggregation in the ITC cell, it was necessary to perform the titration at 400 mM NaCl. Increasing the salt concentration had only a slight effect on 2A binding as judged by EMSA (Figure 4A) and under these conditions we observed a K_D_ of 67 ± 7 nM and a ∼ 1:1 molar ratio of protein to RNA (Figures 4B and C). The large contribution of ΔH (−10.7 ± 0.13 kcal/mol) term to the overall free energy of binding (−9.8 kcal/mol) is consistent with an electrostatic interaction mechanism. To confirm the stoichiometry in solution, we performed a SEC-MALS analysis of the mixture retrieved from the ITC cell (Figure 4D). As expected, the two peaks correspond to an approximate 1:1 RNA-protein complex (early; M_w_ 33,570 Da observed vs. 27,105 Da expected) and excess free RNA (late; M_w_ 10,980 Da observed vs 11,164 Da expected).

**Figure 4.**
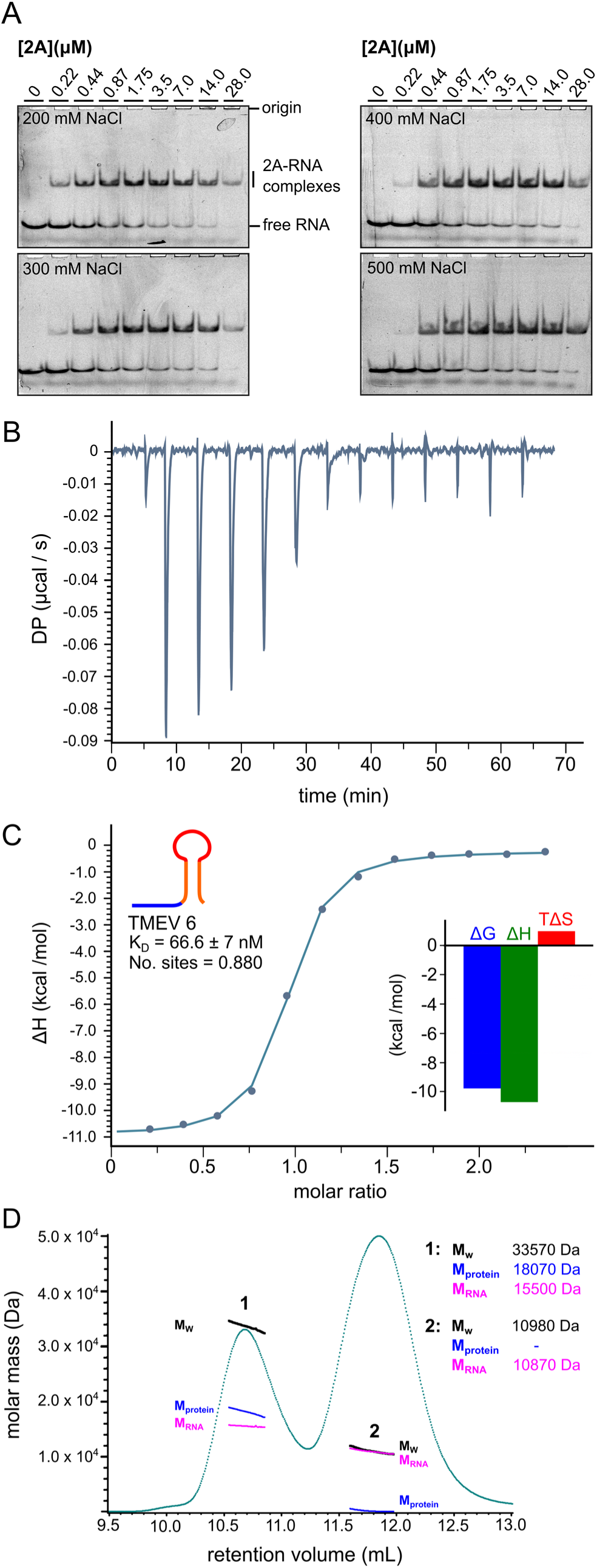
2A binds to the RNA stimulatory element with equimolar stoichiometry and nanomolar affinity. **A.** EMSA analysis showing the effects of salt concentration (indicated) on RNA binding. Experiments were conducted with 50 nM Cy5-labelled TMEV 6 and 2A concentrations between zero and 28 μM. Following non-denaturing electrophoresis, fluorescence was imaged using a Typhoon scanner. **B.** ITC isotherm for a titration of TMEV 6 RNA into 2A protein in the presence of 400 mM NaCl. **C.** Binding curve from titration in B, showing approximately 1:1 molar ratio and nanomolar affinity <Inset> Histogram showing relative contributions of ΔH and TΔS terms to the overall exergonic interaction. **D.** SEC-MALS analysis of 2A-TMEV 6 RNA complex in buffer containing 200 mM NaCl. The 280 nm UV absorbance trace is shown (green). Weight-averaged molar masses across the indicated peaks are listed, along with mass contributions from the protein (blue) and RNA (pink) components, following a protein conjugate analysis. The two peaks correspond to the RNA-protein complex (peak 1; M_w_ 33.6 kDa) and the excess RNA (peak 2; M_w_ 11.0 kDa).

### Snapshots of translation in TMEV-infected cells by ribosome profiling

We extended our examination of 2A-stimulated PRF by analysing TMEV-infected cells using ribosome profiling, a deep sequencing-based technique that gives a global snapshot of the positions of translating ribosomes at sub-codon resolution (31, 32). In addition to WT virus, two mutant viruses were employed. Virus SS has two synonymous mutations in the slippery sequence (G_GUU_UUU to A_GUG_UUU) that prevent frameshifting (23, 24). Virus M3 has the WT slippery sequence, but contains the M3 mutations in the 2A protein described above (Figure 1D), rendering it unable to bind to the PRF-stimulatory RNA stem-loop (23).

BSR cells, a single cell clone of the *Mesocricetus auratus* BHK-21 cell line, were selected for the ribosome profiling due to their relative genetic homogeneity. Thus, we first verified in BSR cells several features of TMEV infection observed in BHK-21 cells (11, 23). The small-plaque phenotype previously observed for the SS and M3 mutant viruses (22–24) was confirmed (Figure S5A), and expression of 2A was detectable from eight hours post-infection (hpi) onwards. Levels increased up to 12 hpi, at which point cytopathic effects become fairly extensive (Figure S5B).

Metabolic labelling experiments were carried out to verify the occurrence of highly efficient frameshifting in TMEV-infected BSR cells. Viral proteins downstream of the frameshift site are only translated by ribosomes which have not undergone frameshifting, as those that frameshift encounter a stop codon in the −1 frame eight codons downstream of the shift site. Frameshift efficiency was estimated from the ratio of downstream to upstream products (Figure 5A, Figure S5C), normalised by the frameshift-defective M3 mutant as a control (detailed in Methods, and previously described in refs. 11,33). The mean WT frameshift efficiency was found to increase from 63% at 10 hpi to 78% at 12 hpi (Figure 5B, blue bars) (two-tailed Welch’s *t*-test: *t* = −2.92, *df* = 2.80, *p* = 0.067) similar to the 74 – 82% range previously calculated by metabolic labelling of infected BHK-21 cells (11, 23). The frameshift efficiency of the SS mutant virus was low at both timepoints (5% and 1% at 10 and 12 hpi respectively), as previously seen for TMEV (23) and EMCV (24).

**Figure 5.**
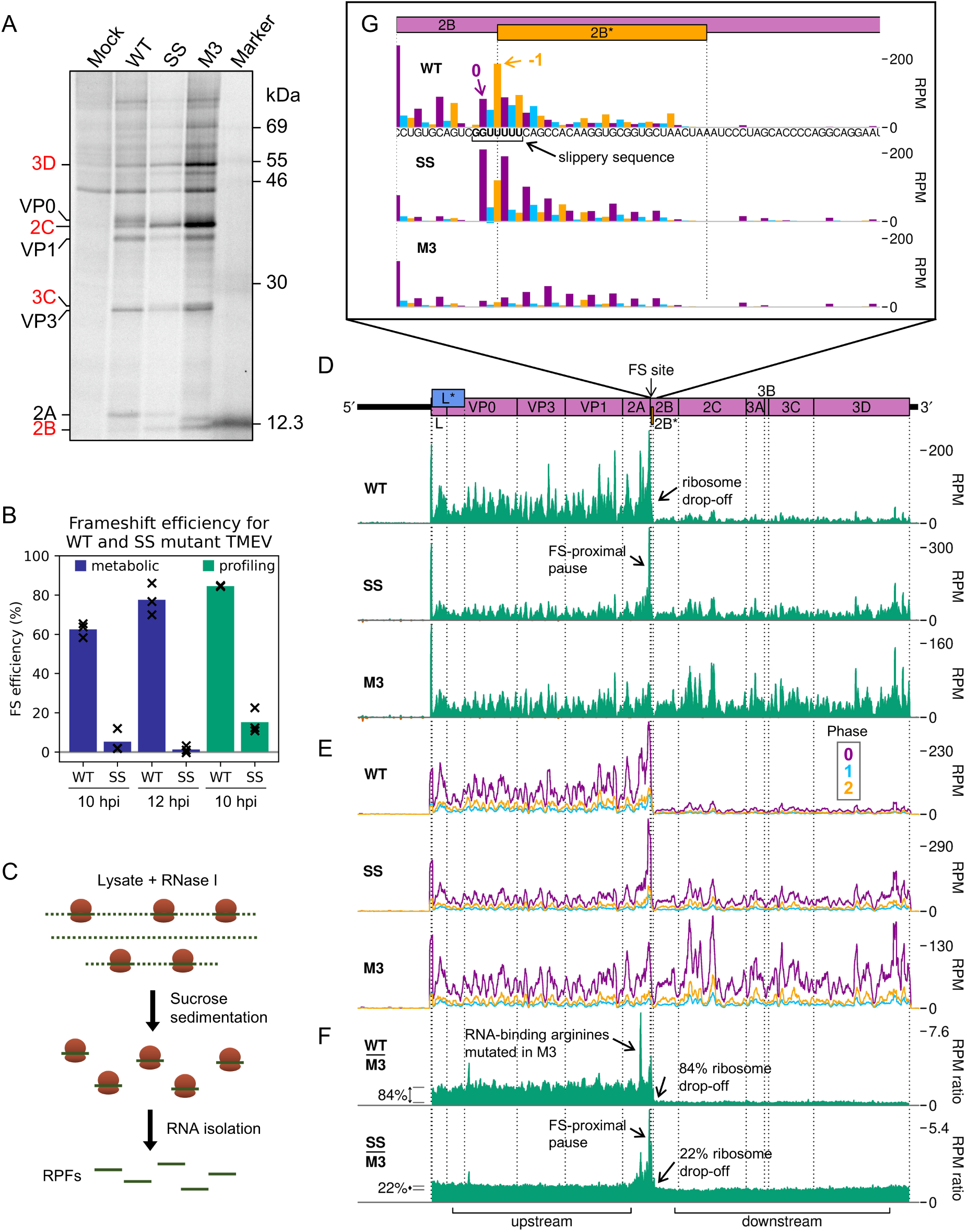
2A stimulates -1 PRF with 85% efficiency and leads to ribosomal pausing in infected cells. **A.** Metabolic labelling of BSR cells infected with WT, SS, or M3 TMEV and harvested at 10 hpi. Positions of TMEV proteins are indicated, with those downstream of the frameshift site written in red. **B.** Frameshift (FS) efficiency in infected cells, calculated from metabolic labelling (blue bars) and ribosome profiling (green bars). Bars represent the mean of three replicates, with values for each replicate indicated by crosses. **C.** Schematic of ribosome profiling methodology. Cells are flash frozen before lysis. RNase I is added to digest regions of unprotected RNA, then ribosomes and enclosed ribosome-protected fragments (RPFs) of RNA are purified. RPFs are released and prepared for high-throughput sequencing to determine the positions of translating ribosomes at the time of cell lysis. **D.** RPF densities in reads per million mapped reads (RPM) on WT, SS and M3 viral genomes (top panel), after application of a 15 nt running mean filter, from cells harvested at 10 hpi. Positive sense reads are plotted in green (above the horizontal axis), negative sense in red (below the horizontal axis; negligible amounts). In all plots, RPF densities from replicate 3 are shown, plotted at the inferred position of the ribosomal P site. **E.** Positive-sense RPF densities from D, coloured according to phase (purple, blue, and yellow represent RPFs whose 5′ ends map to, respectively, the 1st, 2nd or 3rd nucleotides of polyprotein-frame codons, defined as phases 0, +1/−2, and +2/−1), after application of a 15 codon running mean filter. Frames for each viral ORF are designated with respect to the polyprotein (set to 0) and indicated on the genome map by colour and by an offset on the y axis in the corresponding direction. **F.** Ratio of WT or SS RPF density at each position on the genome relative to the RPF density on the M3 mutant genome. UTRs were excluded, only positive sense reads were used, and a 15 nt running mean filter was applied before the division. Regions defined as “upstream” and “downstream” in the ribosome profiling frameshift efficiency calculations are annotated below, and average densities for these regions displayed to the left of the plots with grey lines. **G.** Inferred positions of ribosomal P sites at the slippery sequence and in the 2B* region, coloured according to the phase of RPF 5′ ends as in E. The genome sequence in this region is underlaid beneath the data in the top panel, and all libraries are set to the same scale on the y axis, with no running mean filter applied.

The 10 hpi timepoint was selected for the ribosome profiling. BSR cells were infected with TMEV WT, SS, or M3, or mock-infected, and harvested by snap-freezing in liquid nitrogen. Cells were lysed and the presence of 2A verified by western blotting (Figure S6A). Lysates were treated with RNase I and ribosome-protected mRNA fragments (RPFs) were harvested by pelleting the ribosomes through sucrose with subsequent phenol extraction (Figure 5C). RPFs were ligated to adapters, cloned, and deep sequenced. Reads were mapped to the host (*M. auratus*) and viral (based on NC_001366) genomes (Table S2) to determine precisely the locations of translating ribosomes. Quality control analysis, carried out as previously described (34), indicated that the datasets were of high quality (Figure S6). RPFs mapping to coding sequences exhibited the characteristic length distribution with peaks at 20 – 22 nt and 28 – 30 nt (Figure S6B), corresponding to the two distinct lengths of mRNA protected from nuclease digestion by the translating ribosome (31,35–38). In mammalian cell lysates treated with sufficient nuclease so that all unprotected regions of mRNA are fully digested (“trimmed”), there is a distance of ∼ 12 nt between the 5′ end of the RPF and the first nucleotide in the P site of the ribosome (31). This can be used to infer the frame of translation. In our libraries, the majority of CDS-mapping RPF 5′ ends map to the first nucleotide position of codons (herein termed phase 0) (Figure S6C), indicating successful nuclease trimming. RPFs map to coding sequences with a triplet periodicity reflective of the length of a codon and, as expected, few RPFs map to the UTRs (Figure S6D).

### 2A-stimulated frameshifting occurs with 85% efficiency in infected cells

To analyse translation of the viral RNA, we plotted RPF distribution on the viral genome (Figure 5D-G, Figure S7). There is a clear dominance of the 0 phase throughout the genome (Figure 5E), including within the +1-frame L* ORF which overlaps L and VP0, indicating that the L* ORF is not highly translated. Ribosomes that undergo frameshifting at the 2B* shift site translate only eight codons in the −1 frame before encountering a termination codon. This results in a striking drop-off in ribosome density on the WT genome (Figure 5D). No such decrease in density is seen after the frameshift site on the SS virus genome, and there is actually a slight increase in read density in this region on the M3 genome. This could be due to differences between the two regions in mean translation rate or the extent of various biasing effects inherent in the ribosome profiling procedure, such as ligation, PCR or nuclease biases (39). In order to control for these effects and highlight translational features related to the presence of a functional 2A and/or shift site, the read densities on the WT and SS genomes were divided by those in the corresponding position on the M3 genome (Figure 5F, Figure S7B). Frameshift efficiency was calculated from the ratio of M3-normalised RPF density in the regions downstream and upstream of the frameshift site, revealing a mean WT frameshift efficiency of 85%, which to our knowledge is the highest −1 PRF efficiency thus far recorded in any natural system (11,22,23) (Figure 5B**, green bars**). This is significantly greater than the 63% measured by metabolic labelling at the same timepoint (two-tailed Welch’s *t*-test: *t* = −10.2, *df* = 2.03, *p* = 0.0090). The profiling assay (with M3 normalisation) is expected to be substantially more accurate than the metabolic labelling approach, which suffers from lower sensitivity (densitometry of a few protein bands versus high throughput sequencing) and lower temporal resolution (1 hour labelling vs snap-freezing translating ribosomes).

The occurrence of a highly efficient frameshift in WT TMEV was verified by the observation of a marked shift in the dominant RPF phase from 0 in the upstream region, to −1/+2 in the 2B* transframe region (Figure S8A). It should be noted that in frameshift efficiency calculations, neither the profiling nor the metabolic labelling experiments would reliably distinguish between ribosome drop-off due to termination at the 2B* stop codon post-frameshifting and ribosome drop-off at the StopGo site located just five codons upstream. However, StopGo is generally very efficient in cultured cells, with little drop-off (40, 41), and this is evident here, as no obvious decrease in RPF density occurs after the StopGo motif in the M3 mutant (Figure 5D, Figure S7A). Surprisingly, the mean frameshift efficiency of the SS mutant was high, at 15%. Similar residual frameshift activity has been observed at other highly efficient frameshift signals with analogous mutations designed to knock out frameshifting (42, 43). This may be reflective of the very strong frameshift-stimulatory activity of the 2A-RNA complex facilitating frameshifting even despite unfavourable codon-anticodon repairing in the −1 frame. However, prolonged pausing of ribosomes over the mutant slippery sequence (discussed below) could potentially influence mechanisms other than frameshifting that may contribute to this observed drop-off, such as ribosome rescue pathways (44).

### Profiling reveals multiple 2A-related ribosomal pausing events

Ribosome pausing over the slippery sequence has long been considered a mechanistically important feature of PRF (16-18,21). However, while observed to a small extent on WT shift sites *in vitro* (23,42,45–47), it has been elusive in ribosome profiling data (34, 48). However, if the slippery sequence is mutated to prevent frameshifting, a measurable pause is seen both *in vitro* (18,23,27) and in profiling experiments (22), perhaps reflecting a reduced ability of non-frameshifting ribosomes to resolve the topological problem posed by the downstream frameshift-stimulatory element. Ribosome profiling can allow the identification of ribosome pauses with single-nucleotide precision, but at the level of individual nucleotides, the profiles can be strongly affected by nuclease, ligation and potentially, other biases introduced during library preparation. However, by comparing the WT and SS genome ribosome profiles with the M3 genome ribosome profile, many of these biases are normalised and allow changes in dwell time to be identified that are likely to arise as a result of 2A binding.

Apart from ribosome drop-off at the 2B* stop codon, the region of greatest difference in a comparison of the WT and M3 profiling datasets was surprisingly not at the frameshift site, but in the middle of the 2A ORF (Figure 5F, Figure S7B). This pause likely corresponds to ribosomes translating the RNA-binding arginine residues (R85 and R87), which are mutated to alanine in the M3 genome. The pause on the M3-normalised plot likely reflects increased decoding time for the arginine-encoding CGC codons in the WT, which are relatively poorly adapted to the cellular tRNA pool compared to the alanine-encoding GCC codons in the M3 mutant (49, 50) (Figure S8B and C). This peak is present to a lesser extent on the SS genome, potentially due to the slightly slower replication kinetics of this mutant virus (11) meaning there are fewer copies of the viral genome to deplete the cellular supply of the relevant amino-acylated tRNA.

Looking specifically at the frameshift site, a single-nucleotide resolution plot of reads mapping to this region reveals a peak on the SS mutant genome corresponding to a ribosome paused with the GUG codon of the mutated slippery sequence (corresponding to WT GUU) in the P site (Figure 5G, Figure S7C). This putative pause is present to a lesser extent on the WT genome (marked 0 in purple), but not the M3 genome, indicating it is related to the presence of functional 2A. The UUU codon of the slippery sequence also appears to have a much larger phase 0 peak in SS than M3, however this may be enhanced by potential “run-on” effects in which a fraction of ribosome pauses may be able to resolve during cell harvesting and ribosomes translocate to the next codon (50). On both WT and SS genomes, a noticeable peak is present two nucleotides downstream of the main slippery sequence pause, corresponding to ribosomes which have frameshifted and then translocated one codon, and further peaks suggestive of −1-frame translation throughout the 2B* ORF, especially on the WT template. Closer inspection of the whole-genome (Figure 5D, Figure S7A) and M3-normalised plots (Figure 5F, Figure S7B) reveals that the pause at the frameshift site actually extends a little further upstream, ending just upstream of the 2A-2B junction formed by the StopGo motif. Further investigation of this region (Figure S8B) reveals prominent pauses in the SS mutant, and to a lesser extent the WT, with ribosomal P sites corresponding to the glutamic acid and methionine residues of the D(V/I)**Ex**NPG|P StopGo motif (pause sites highlighted in bold, where x is methionine in TMEV). These pauses are larger than the pause over the slippery sequence itself. Ribosomal pausing over StopGo motifs has been observed *in vitro* (41) but have been shown to occur with the ribosomal P site corresponding to the conserved glycine residue directly before the separation site (41, 51), suggesting that the pauses we see have another origin.

### Disome profiling provides evidence for ribosome queuing at the frameshift and StopGo sites

Noticing that the main pause over the StopGo motif was nine codons upstream of the pause over the slippery sequence, we wondered whether this might be consistent with the transient formation of disomes, in which the leading ribosome is paused over the slippery sequence. Disomes are routinely excluded during preparation of ribosome profiling libraries by the inclusion of a size-selection step (in this study, 19-34 nt) which selects for monosome-protected fragments (31). For mock, WT-and M3-infected lysates from replicate 3 we carried out two parallel size selection steps, in which the 19-34 nt “monosome” fraction and a 35-65 nt “broad spectrum” fraction were isolated from the same lysate. The length distribution of broad spectrum reads demonstrated local peaks at read lengths of around 51, 54 and 59-63 nt, consistent with expected lengths of RNA protected by disomes (36,52,53) (Figure 6A). Reads of lengths 51-52, 54-55 and 57-64 nt showed a bias in phase composition towards phase 0, indicating a portion of genuine ribosome footprints (Figure 6B). These read lengths were selected for analysis as potential “disome-protected fragments”, and their density plotted on the viral genome at the inferred P site position of the upstream, colliding ribosome (Figure 6C and D).

**Figure 6.**
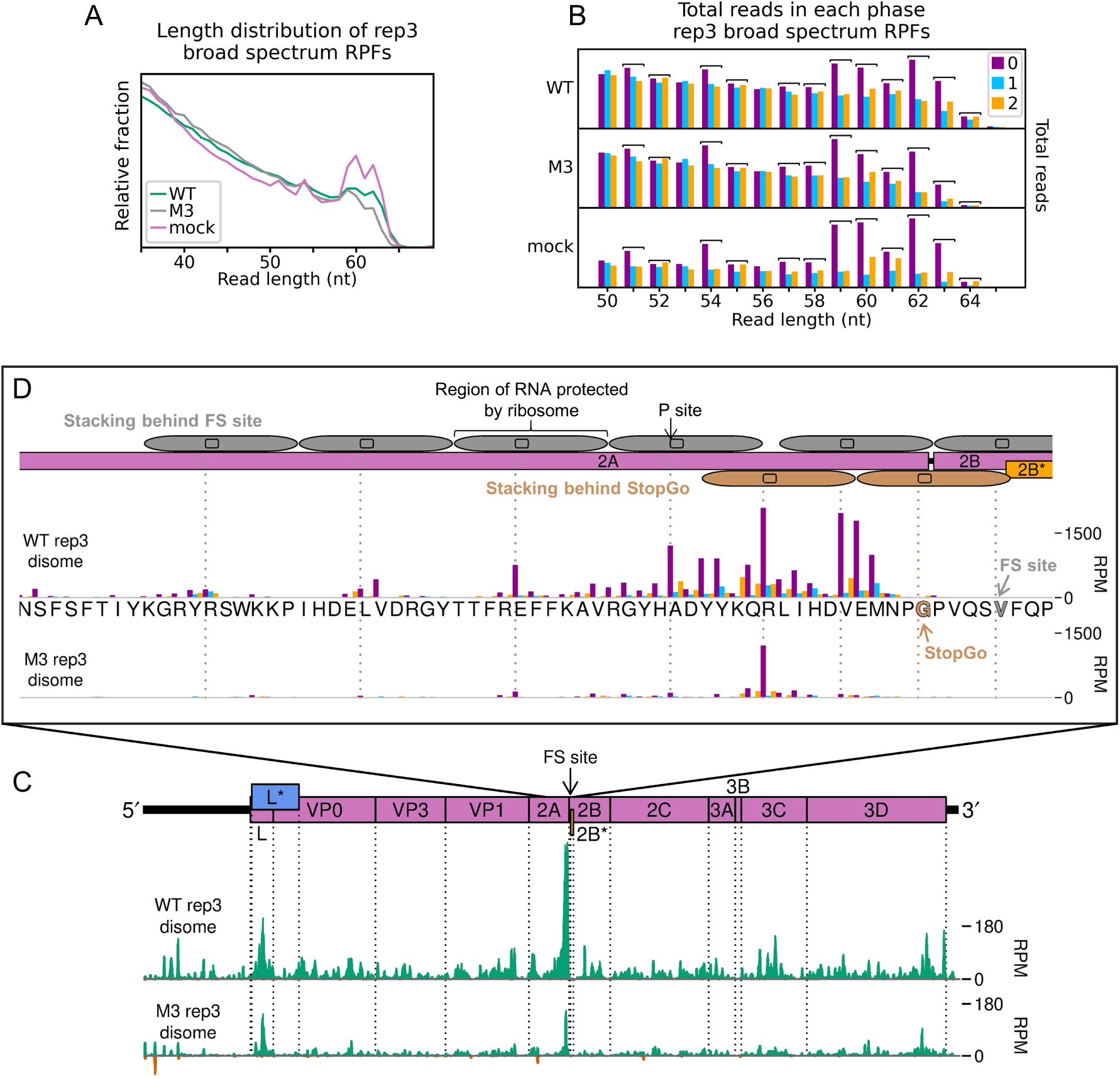
Profiling of disome-protected fragments reveals ribosome stacking at the frameshift site. **A.** Length distribution of reads mapping within host CDSs for broad spectrum libraries. **B.** Total number of reads attributed to each phase, for reads of each specified length, in the broad-spectrum libraries. Read lengths which were selected for inclusion in the “disome-protected fragment” plots (C and D) are indicated by square brackets, and correspond to peaks in A. **C.** Density of disome-protected fragments on the WT and M3 viral genomes, derived from the same lysate as the monosome-protected fragment data plotted in Figure 5 D-G. Reads of lengths 51-52, 54-55 and 57-64 nt were selected for inclusion, and read densities are plotted as RPM after application of a 15 nt running mean filter. Positive sense reads are plotted in green (above the horizontal axis), negative sense in red (below the horizontal axis). In all disome plots, reads are plotted at the inferred P site position of the colliding ribosome. **D.** Density of disome-protected fragments, plotted at inferred P site positions of colliding ribosomes involved in disomes upstream of the 2B* frameshift site, coloured according to phase as in B. The encoded amino acid sequence in this region is underlaid beneath the data in the top panel, and both libraries are set to the same scale on the y axis, with no running mean filter applied. Codons on which leading ribosomes would be expected to pause due to StopGo and frameshifting (FS) are indicated in brown and grey respectively. Positions of ribosomes potentially involved in queue formation behind these pause sites are indicated (FS: above genome map; StopGo: below genome map), with P site positions annotated as dashed vertical lines, in the corresponding colours.

A very prominent peak is visible on the WT genome over the StopGo motif (Figure 6C), and closer inspection reveals that one of the highest peaks in this region is over the valine of the StopGo motif, ten codons upstream of the slippery sequence pause (Figure 6D). This is the expected distance between the P sites of ribosomes involved in a disome (52–54), and would be consistent with disome formation due to a ribosome translating the StopGo motif colliding with a ribosome paused over the slippery sequence. Further, this approximate ten-codon periodicity in distances between peaks extends upstream, consistent with potential formation of ribosome queues up to six ribosomes long (53, 55) (Figure 6D, grey oblongs). An additional peak is evident over the arginine codon ten codons upstream of the StopGo release site, which would correspond to a disome in which the leading ribosome was paused over the conserved glycine codon of the StopGo motif (Figure 6D, brown oblongs). This is evidently a feature of the StopGo site itself and unrelated to the binding of 2A downstream, as it occurs in the M3 dataset as well as the WT. It should be noted that ribosome profiling generates an averaged result of ribosome positions over multiple copies of the viral genome, and formation of the two putative disomes proposed here could not occur simultaneously on a single RNA.

The potential formation of ribosome queues behind the frameshift/StopGo site is supported by the monosome data, in which RPF density gradually increases throughout 2A, reaching a maximum over the StopGo motif. This is particularly apparent in the data from the SS mutant (Figure 5D and F), consistent with the idea that greater pausing over the mutant slippery sequence may be increasing disome formation, pushing the lengths of protected fragments into the disome fraction, and reducing their visibility in the monosome dataset. This is unusual, as no prominent disome peaks were detected near the frameshift site by broad spectrum ribosome profiling of murine coronavirus (34), nor at the −1 PRF site of L-A virus in yeast (56), although we note that −1 PRF is less efficient in both these systems when compared with TMEV. Indeed, the potential formation of disomes at the frameshift site in TMEV represents the first evidence of ribosomes forming queues at a −1 PRF signal in a eukaryotic system. Disome formation has recently been mechanistically implicated in +1 PRF in yeast and +1 and −1 PRF in bacteria (56–59). It may be that the impact of ribosomes colliding at the TMEV PRF site contributes to the complex energetic and conformational landscape required to overcome the translation blockade and break the triplet codon periodicity.

## Conclusion

Our combination of structural, biophysical, biochemical and deep sequencing approaches affords a view of unparalleled detail into the molecular mechanisms underpinning *Theilovirus* protein-stimulated frameshifting. Despite highly divergent sequences within cardioviruses, the TMEV 2A protein adopts the second known occurrence of the beta-shell fold. Whilst the distribution of positively charged residues comprising the putative RNA-binding surface is different from that found in EMCV, it nevertheless recapitulates the same exquisite conformational selectivity for binding to its viral RNA target. Strikingly, we demonstrate that this is a pseudoknot that is likely to exist in equilibrium with the stem-loop previously suggested by structure probing experiments. Despite being 11 nt shorter, the stimulatory RNA element is able to adopt a conformation topologically equivalent to that previously seen in EMCV, involving interactions between the loop and the 5′ extension. In infected cells, stabilisation of this conformation by high-affinity 2A binding represents the ‘switch’ that controls frameshifting and thereby reprogramming of viral gene expression. Our ribosome profiling data reveals that, when invoked, this is up to 85% efficient, representing the highest known −1 PRF efficiency to date. Whilst frameshift-associated pausing is not normally detectable at WT shift sites in profiling data, we show that it is detectable here by analysing RPFs corresponding to disomes, not only at the PRF site but also the adjacent StopGo motif. This is consistent with the relatively long pauses accompanying TMEV frameshifting *in vitro*, and suggests that ribosome collisions are more common than previously thought during translation of the TMEV genome. Taken together, these results suggest that there is a fine balance between necessary ribosome pausing associated with recoding events, and the detrimental effect on viral fitness that may result from these pauses lasting long enough to trigger ribosome quality control pathways, and the degradation of the viral RNA. In future, structural characterisation of the 2A-RNA complex will yield further insights into the molecular mechanisms underlying the potency of this elongation blockade.

## Materials and Methods

### Protein expression and purification

TMEV 2A cDNA was amplified by PCR from plasmid pGEX6P2-based constructs (23) (F 5′ AATTCATATGAATCCCGCTTCTCTCTACCGC 3′; R 5′ AATTGGATCCTTATTAGCCTGGGTTCATTTCTACATC 3′) and cloned into pOPT3G (60) to introduce a 3C protease-cleavable N-terminal GST tag. Recombinant protein was produced in *E. coli* BL21(DE3) pLysS cells. Cultures were grown in 2xTY broth supplemented with 100 μg/mL ampicillin (37 °C, 210 rpm). Expression was induced at A_600_ of ∼ 1 with 1.0 mM isopropyl β-D-1-thiogalactopyranoside (IPTG) and continued overnight (210 rpm, 21 °C, 16h). Bacteria were pelleted (4,000 × g, 4 °C, 20 min), washed in cold PBS and stored at −20 °C. Cells from four litres of culture were thawed in 200 mL lysis buffer (50 mM Tris (HCl) pH 7.4, 500 mM NaCl, 0.5 mM MgCl_2_, 5.0 mM DTT, 0.05% w/v Tween-20 supplemented with 50 μg/mL DNase I and EDTA-free protease inhibitors) and lysed using a cell disruptor (24 kPSI, 4 °C). The insoluble fraction was pelleted by centrifugation (39,000 × g, 40 min, 4 °C) and discarded. Supernatant was incubated (2 h, 4 °C) with 4.0 mL of Glutathione Sepharose 4B resin (GE Healthcare) that had been pre-equilibrated in the same buffer. Protein-bound resin was washed three times by centrifugation (600 × g, 10 min, 4 °C) and re-suspension in 150 mL wash buffer (50 mM Tris (HCl) pH 7.4, 500 mM NaCl, 5.0 mM DTT). Washed resin was transferred to a gravity column and protein was eluted in batch with 20 mL wash buffer supplemented with 25 mM reduced glutathione (1 h, 4 °C). GST-tagged 3C protease was added to the eluate (10 μg/mL) and the mixture was dialysed (3K molecular weight cut-off (MWCO), 4 °C, 16 h) against 2 L wash buffer to remove the glutathione. Dialysed proteins were then re-incubated with Glutathione Sepharose 4B resin (as above; 2 h, 4 °C) to remove the cleaved GST and GST-3C protease. The flow-through was subjected to heparin-affinity chromatography to remove nucleic acids. Samples were loaded on a 10 mL HiTrap Heparin column (GE Healthcare) at 2.0 mL/min, washed with two column volumes of buffer A (50 mM Tris (HCl) pH 7.4, 500 mM NaCl, 1.0 mM DTT and eluted with a 0% → 100% gradient of buffer B (50 mM Tris (HCl) pH 8.0, 1.0 M NaCl, 1.0 mM DTT) over 20 column volumes. After removal of nucleic acids, protein became aggregation-prone and precipitated at low temperatures, therefore all subsequent steps were performed at 20 °C. Fractions corresponding to the 2A peak were pooled and concentrated using an Amicon® Ultra centrifugal filter unit (10K MWCO, 4,000 × g) prior to size exclusion chromatography (Superdex 75 16/600 column; 10 mM HEPES pH 7.9, 1.0 M NaCl, 1.0 mM DTT). Purity was assessed by 4-20% gradient SDS-PAGE, and protein identity verified by mass spectrometry. Purified protein was used immediately for crystallisation trials or was concentrated (∼ 4.4 mg/mL, 282 μM), snap-frozen in liquid nitrogen and stored at −80 °C.

### Size-exclusion chromatography coupled to multi-angle laser scattering (SEC-MALS)

For studies of the TMEV 2A protein in isolation, a Superdex 75 increase 10/300 GL column (GE Healthcare) was equilibrated with 20 mM Tris (HCl) pH 7.5, 1.0 M NaCl (0.4 mL/min flow, 25 °C). Per experiment, 100 μL of protein was injected at concentrations of 3.1, 0.8 and 0.38 mg/mL (molar concentrations of 200, 51.5 and 24.5 μM, respectively). The static light scattering, differential refractive index, and the UV absorbance at 280 nm were measured by DAWN 8+ (Wyatt Technology), Optilab T-rEX (Wyatt Technology), and Agilent 1260 UV (Agilent Technologies) detectors. The corresponding molar mass from each elution peak was calculated using ASTRA 6 software (Wyatt Technology) using the differential refractive index and a dn/dc value of 0.186 to calculate the protein concentration. For studies of 2A-RNA complexes, samples were recovered directly from the ITC cell after confirmation of binding and concentrated to an A_280_ of ∼ 3.1 prior to injection of 100 uL onto a Superdex 75 increase 10/300 GL column pre-equilibrated in 50 mM Tris (HCl) pH 7.4, 200 mM NaCl (0.4 mL/min flow, 25 °C). Data were recorded as above, and to estimate the relative contributions of protein and RNA to molar mass, a protein conjugate analysis was performed within ASTRA 6, using a protein dn/dc value of 0.186 and an RNA (modifier) dn/dc value of 0.168. Prior to this analysis, extinction coefficients (at 280 nm) were determined experimentally from protein-only and RNA-only peaks using the “UV extinction from RI peak” method in ASTRA 6.

### Protein crystallization

Purified TMEV 2A was concentrated to 4.38 mg/mL in 50 mM Tris (HCl) pH 7.4, 1.1 M NaCl, 1.0 mM DTT. Sitting-drop vapor diffusion experiments were set up in 96-well MRC plates with 80 μL reservoir solution, 200 nL protein and 200 nL crystallization buffer. Diffraction-quality crystals grew in 0.2 M KBr, 0.2 M potassium thiocyanate, 0.1 M sodium cacodylate pH 6.5, 3% w/v poly-γ-glutamic acid 200-400 and 10% w/v PEG-MME 2000. Crystals were harvested in nylon loops and cryo-protected by removal from the mother liquor through a 0.5 μL layer of crystallization buffer that had been supplemented with 20% v/v glycerol, prior to flash-cooling by plunging into liquid nitrogen.

### X-ray data collection, structure determination, refinement and analysis

Datasets of 900 images (Table S1) were recorded from two crystals at beamline I04-1, Diamond Light Source on a Pilatus 6M detector (Dectris), using 47% transmission, an oscillation range of 0.2 ° and an exposure time of 0.2 s per image. Data were collected at a wavelength of 0.9159 Å and a temperature of 100 K. Reflections were indexed and integrated with DIALS (61) (highest resolution crystal) or XDS (62) (structure determination crystal) and data were scaled and merged with AIMLESS (63) within the Xia2 data reduction pipeline (64). Resolution cut-off was decided by a CC_1/2_ value ≥ 0.5 and an I/σ(I) ≥ 2.0 in the highest resolution shell (65). The structure was solved by single-wavelength anomalous dispersion (SAD) analysis of the structure determination crystal, using anomalous signal from bromine present in the crystallisation buffer. SAD phasing was performed using autoSHARP (66), implementing SHELXD for substructure determination (67), SHARP for density modification (66) and ARP/wARP (68) for automated model building. This placed 123 residues out of 133 (92%) in the single chain that comprised the asymmetric unit of the cubic *P*2_1_3 cell. This preliminary model was subsequently used as a molecular replacement search model to solve a higher resolution dataset (highest resolution crystal) using Phaser (69). This model was then subjected to several rounds of manual adjustment using COOT (70) and refinement with phenix.refine (71). Upon completion of model building, ISOLDE (72) was used to improve model geometry and resolve clashes prior to a final round of refinement using phenix.refine. MolProbity (73) was used to assess model geometry including Ramachandran outliers, bad rotamers and mainchain geometry deviations throughout the refinement process. For the electrostatic potential calculations, PDB2PQR (74) and PROPKA were used to estimate protein pKa values and assign partial charges. Electrostatic surface potentials were calculated using APBS (75). Relative solvent-accessible surface areas per residue were calculated using GetArea (76) and crystallographic interaction interfaces were assessed using PDBePISA (77). Structural figures depicting crystallographic data (cartoon, stick and surface representations) were rendered in PyMOL (Schrödinger LLC). The representation of surface conservation was generated using ConSurf (78).

### Nucleotide and protein sequence alignments

The Logo plot of nucleotide sequence conservation at the PRF region was generated from a selection of divergent isolates (*Cardiovirus A* – M8161, M22457, KP892662, LC585221, KC310737, JX257003; *Cardiovirus B* – EU542581, M20301, MF172923, MF352420; *Cardiovirus D* – EU376934; *Cardiovirus E, F* and unassigned – KY432928, KY432930, KY855434, KF823815) with WebLogo 2.8.2 (79) using the default parameters. For 2A protein sequence alignments, the match > align tool in UCSF Chimera (80) was first used to generate a seed alignment based on superposed structures of EMCV and TMEV 2A, prior to the subsequent alignment of other selected divergent TMEV-like (MF352420, M20301, MF172923, EU542581, EU376394, KY432930, KY432928, KF823815, KY855434) and EMCV-like (LC585221, KP892662, M81861, M22457, JX257003, KC310737) 2A sequences. JalView (81) was used to visualise the alignment, calculate the consensus sequence and generate the associated Logo plot.

### Electrophoretic Mobility Shift Assay (EMSA)

RNA oligonucleotides (IDT) were reconstituted in distilled water. A 5′ Cy5 fluorescent label was incorporated using the 5′ EndTag kit (Vector Labs) according to the manufacturer’s instructions, prior to phenol:chloroform extraction, ethanol precipitation and aqueous resuspension. Binding reactions of 10 μL contained 1.0 μL 500 nM Cy5-RNA, 1.0 μL TMEV 2A at concentrations of 280, 140, 70, 35, 17.5, 8.7, 4.4, and 2.2 μM in 10 mM HEPES pH 7.9, 1.0 M NaCl, 5.0 μL 2 × buffer (20 mM Tris (HCl) pH 7.4, 80 mM NaCl, 4.0 mM magnesium acetate, 2.0 mM DTT, 10% v/v glycerol, 0.02% w/v bromophenol blue, 200 μg/mL porcine liver tRNA and 800 U /mL SUPERase-In [Invitrogen]) and 3.0 μL distilled water. Final concentrations in the binding reactions were therefore 50 nM RNA, 1 × buffer, ∼ 140 mM NaCl and TMEV 2A at 28.0, 14.0, 7.0, 3.5, 1.75, 0.87, 0.44 and 0.22 μM. All binding reactions were prepared on ice, and samples were incubated at 37 °C for 20 min before analysis by non-denaturing 10% acrylamide/TBE PAGE (25 min, 200 V constant). Gels were imaged with a Typhoon FLA-7000 (GE) using the 635 nm laser / R670 filter.

### Microscale Thermophoresis (MST)

Synthetic RNA oligonucleotides (IDT) were labelled at the 5′ end with Cy5 as described above, prior to purification using Clean and Concentrator kit (Zymo). TMEV 2A in 10 mM HEPES pH 7.9, 1.0 M NaCl was diluted in 2 × buffer (20 mM Tris (HCl) pH 7.4, 80 mM NaCl, 4.0 mM magnesium acetate, 2.0 mM DTT) to a final concentration of 20 μM. For the measurement, a series of 16 1:1 dilutions was prepared and each ligand dilution was mixed with one volume of labeled TMEV RNA. Final concentrations in the binding reactions were therefore 5.0 nM RNA, 1 × buffer, ∼ 140 mM NaCl and TMEV 2A ranging from 0.0006 to 20 μM. The reaction was mixed and the samples loaded into Monolith NT.115 Premium Capillaries (NanoTemper Technologies). Measurements were performed using a Monolith NT.115Pico instrument (NanoTemper Technologies) at an ambient temperature of 25 °C. Instrument parameters were adjusted to 5% LED power and medium MST power. Data of two independently pipetted measurements were analysed for fraction bound using initial fluorescence (MO.Affinity Analysis software, NanoTemper Technologies). The non-binding RNAs (namely TMEV 2 and were normalized using Prism 8.0.2 (GraphPad). Data was plotted using Prism 8.0.2 (GraphPad) software.

### Isothermal Titration Calorimetry (ITC)

ITC analyses were carried out at 25 °C using a MicroCal PEAQ-ITC (Malvern Panalytical). RNAs and proteins were dialysed (24 h, 25 °C) into buffer (50 mM Tris (HCl) pH 7.4, 400 mM NaCl) before performing experiments. Final concentrations of protein and RNA after dialysis were determined by spectrophotometry (A_280_ and A_260_, respectively), using theoretical extinction coefficients based on the primary sequence of each component. RNA (60 μM) was titrated into protein (5 μM) with 1 × 0.4 μL injection followed by 12 × 3.0 μL injections. Control titrations of RNA into buffer, buffer into protein and buffer into buffer were also performed. Data were analyzed using the MicroCal PEAQ-ITC analysis software (Malvern Panalytical) and binding constants determined by fitting a single-site binding model.

### In vitro transcription

For *in vitro* frameshifting assays, a 105 nt portion of the TMEV genome containing the GGUUUUU shift site flanked by 6 nt upstream and 92 nt downstream was cloned into the dual luciferase plasmid pDluc at the XhoI/BglII sites (82). The sequence was inserted between the Renilla and firefly luciferase genes so that firefly luciferase expression is dependent on −1 PRF. Wild-type or derivative frameshift reporter plasmids were linearized with FspI and capped run-off transcripts generated using T7 RNA polymerase as described (83). Messenger RNAs were recovered by phenol/chloroform extraction (1:1 v/v), desalted by centrifugation through a NucAway Spin Column (Ambion) and concentrated by ethanol precipitation. The mRNA was resuspended in water, checked for integrity by agarose gel electrophoresis, and quantified by spectrophotometry.

### In vitro translation

Messenger RNAs were translated in nuclease-treated rabbit reticulocyte lysate (RRL) (Promega). Typical reactions were composed of 90% (v/v) RRL, 20 μM amino acids (lacking methionine) and 0.2 MBq [^35^S]-methionine and programmed with ∼50 μg/ml template mRNA. Reactions were incubated for 1 h at 30 °C. Samples were mixed with 10 volumes of 2× Laemmli’s sample buffer, boiled for 3 min and resolved by SDS-PAGE. Dried gels were exposed to a Storage Phosphor Screen (PerkinElmer), the screen was then scanned in a Typhoon FLA7000 using the phosphor autoradiography mode. Bands were quantified using ImageQuant™TL software. The calculations of frameshifting efficiency (% FS) took into account the differential methionine content of the various products and % FS was calculated as % −1FS = 100 × (IFS/MetFS) / [(IS/MetS) + (IFS/MetFS)]. In the formula, the number of methionines in the stop, − are denoted by MetS, MetFS respectively; while the densitometry values for the same products are denoted by IS and IFS respectively. All frameshift assays were carried out a minimum of three times.

### Cells and viruses

BSR (single cell clone of BHK-21 cells, species *Mesocricetus auratus*, provided by Polly Roy, LSHTM, UK) were maintained in Dulbecco’s modified Eagle’s medium (DMEM), high glucose, supplemented with L-glutamine (1mM), antibiotics, and fetal bovine serum (FBS) (5%), at 37 °C and 5% CO_2_. Cells were verified as mycoplasma-free by PCR (e-Myco *plus* Mycoplasma PCR Detection Kit, iNtRON Biotechnology). Cells were seeded to achieve 80% confluence on the day they were infected with virus stocks listed in (23), based on GDVII isolate NC_001366 with three nucleotide differences present in WT and mutant viruses (G2241A, A2390G and G4437A; nt coordinates with respect to NC_001366). All infections, except for plaque assays, were carried out at a MOI of 3 in serum-free media, or media only for the mock-infected samples. After incubation at 37 °C for 1 h, inoculum was replaced with serum-free media supplemented with FBS (2%), and infected cells were incubated at 37 °C until harvesting or further processing.

### Plaque assays

BSR cells in 6-well plates at 90% confluence were inoculated with serial dilutions of virus stocks for 1 h then overlaid with 2ml DMEM supplemented with L-glutamine (1.0 mM), antibiotics, FBS (2%), and carboxymethyl cellulose (0.6%). Plates were incubated at 37 °C for 48 h then fixed with formal saline and stained with 0.1% toluidine blue.

### Western blots of 2A expression

BSR cells in 35mm dishes were inoculated for 1 h at a MOI of 3, and incubated at 37 °C until 2, 4, 6, 8, 10, or 12 hpi. Cells were washed with cold PBS and lysed in 200µl 1X radioimmunoprecipitation assay (RIPA) buffer (50 mM Tris (HCl) pH 8, 150 mM sodium chloride, 1% NP-40 substitute, 0.5% sodium deoxycholate, 0.1% SDS) supplemented with Halt™ Protease Inhibitor Cocktail (1X). Samples were resolved by 4-20% gradient SDS-PAGE and transferred to 0.2 μm nitrocellulose membranes. Membranes were blocked with 5% w/v milk dissolved in PBS (1 h, 25 °C). Primary antibodies were diluted in 5% w/v milk, PBS, 0.1% v/v Tween-20 and incubated with membranes (1 h, 25 °C). After three washes in PBS, 0.1% v/v Tween-20, membranes were incubated with IRDye fluorescent antibodies in 5% w/v milk, PBS, 0.1% v/v Tween-20 (1 h, 25 °C). Membranes were washed three times in PBS, 0.1% v/v Tween-20 and rinsed in PBS prior to fluorescent imaging with an Odyssey CLx platform (LI-COR). Antibodies used were rabbit polyclonal anti-2A (22) (1/1000); mouse monoclonal anti-GAPDH (1/20,000, clone G8795, Sigma-Aldrich), goat anti-rabbit IRDye 800 CW (1/10,000, LI-COR), and goat anti-mouse IgM (μ chain specific) IRDye 680RD (1/10,000, Li-Cor).

### Metabolic labelling of infected cells

BSR cells in 24-well plates were infected at a MOI of 3 in a volume of 150 μl. After 1 h, the inoculum was replaced with DMEM containing 2% FBS (1 ml) and cells were incubated at 37 °C for 5, 7, or 9 h delineating the 8, 10, or 12 hpi -timepoints respectively. At the set timepoint post-infection, cells were incubated for 1 h in methionine-and serum-free DMEM, then radiolabelled for 1 h with [^35^S]-methionine at 100 μCi ml^−1^ (∼1,100 Ci mmol^−1^) in methionine-free medium. Cells were harvested and washed twice by resuspension in 1 ml of ice-cold phosphate-buffered saline (PBS) and pelleted at 13,000 g for 2 min. Cell pellets were lysed in 40 μl 4 × SDS–PAGE sample buffer and boiled for 5 min before analysis by SDS–PAGE. Dried gels were exposed to X-ray films or to phosphorimager storage screens. Image analysis was carried out using ImageQuantTL 7.0 to quantify the radioactivity in virus-specific products. Bands which were quantifiable for all three viruses (Table S3) were carried forward for normalisation by methionine content and then by an average of the quantifiable proteins upstream of the frameshift site as a loading control. Bands in the WT and SS lanes were normalised by the counterpart bands in the M3 lane to control for differences in protein turnover. Frameshift efficiency (%) is given by the equation 100 × [1-(downstream/upstream)], where downstream and upstream represent the average of the fully normalised intensity values for proteins downstream and upstream of the frameshift site, respectively.

### Ribosome profiling library preparation

BSR cells in 9 cm^2^ dishes were inoculated in triplicate for 1 h at 37 °C at a MOI of 3, or media only for the mock-infected samples. After inoculation, cells were incubated in serum-free media supplemented with FBS (2%) at 37 °C for 9 h. Cells were harvested and ribosome-protected fragments (RPFs) purified and prepared for next-generation sequencing as described (84), based on (32,85,86), with the following modifications. The cycloheximide pre-treatment was omitted from the harvesting protocol and cells were instead washed with warm PBS before snap-freezing in liquid nitrogen. RNase I (Ambion) was added to final concentration 0.5 U/µl (replicate 1) or 2.5 U/µl (replicate 2 and 3 – higher concentration to ameliorate incomplete trimming noticed in replicate 1), and digestion inhibited by adding SUPERase-In RNase inhibitor (Invitrogen) to final concentration 0.13 U/µl and 0.67 U/µl, respectively. The range of fragment sizes selected during polyacrylamide gel purification before ligation to adapters was increased to 19-34 nt, and size ranges for post-ligation gels adjusted accordingly. For replicate 3 samples, two gel slices were excised per library, to purify monosome-protected (19-34 nt) and broad spectrum (35-65 nt) fragments from the same lysate. Depletion of ribosomal RNA was carried out solely by use of the RiboZero Gold Human/Mouse/Rat kit (Illumina). Adapter sequences were based on the TruSeq small RNA sequences (Illumina), with an additional seven random nucleotides at the 5′-end of the 3′-adapter and the 3′-end of the 5′-adapter, to reduce the effects of ligation bias and enable identification of PCR duplicates. For the broad-spectrum libraries only the 3′-adapter contained the 7 random nucleotides. Libraries were deep sequenced on the NextSeq 500 platform (Illumina), and data made publicly available on ArrayExpress under accession numbers E-MTAB-9438 and E-MTAB-9437.

### Ribosome profiling data analysis

Adapter sequences were removed and reads resulting from RNA fragments shorter than 19 nt discarded using fastx_clipper (version 0.0.14, parameters: -Q33 -l 33 -c -n -v). Sequences with no adapters and those which consisted only of adapters were discarded. PCR duplicates were removed using awk, and the seven random nucleotides originating from the adapters were trimmed from each end of the reads using seqtk trimfq (version 1.3). Reads were aligned to reference databases using bowtie1, allowing one mismatch and reporting only the best alignment (version 1.2.3, parameters: -v 1 --best), in the following order: rRNA, virus genome (vRNA), mRNA, ncRNA, mtDNA, and gDNA. Quality control analysis indicated some contamination of replicate 2 libraries with *E. coli* RNA (BL21). These reads do not exhibit the expected features of genuine RPFs, however, indicating that the contamination occurred after lysates were harvested and thus they do not affect our conclusions. Reads were re-mapped with the addition of a BL21 reference database (CP047231.1) before vRNA, to remove these reads. The rRNA database comprised GenBank accession numbers NR_003287.2, NR_023379.1, NR_003285.2 and NR_003286.2. The mRNA database was compiled from the 36827 *M. auratus* GenBank RefSeq mRNAs available on 17^th^ Nov 2017, after removing transcripts with annotated changes in frame. The ncRNA and gDNA databases are from *M. auratus* Ensembl release 90 (genome assembly 1.0). Viral genome sequences were verified by *de novo* assembly with Trinity (version 2.9.1), and reversion rates for mutated bases were verified as below 0.5%. For the broad-spectrum dataset, reads with no detected adapter were not discarded by fastx_clipper, PCR duplicates were not removed, and two mismatches were allowed during bowtie1 mapping, except to BL21 *E. coli*, for which one mismatch was allowed.

To visualise RPF distribution on the viral genome, the number of reads with 5′-ends at each position was counted and divided by the total number of positive-sense vRNA and host mRNA reads for that library, to normalise for library size and calculate reads per million mapped reads (RPM). For plots covering the entire virus genome, a sliding window running mean filter of 15 nt was applied. To generate the plot of WT and SS data normalised by M3 data, UTRs plus a small buffer (7 nt at the 5′-end of the CDS and 27 nt at the 3′-end) were excluded and the result of the 15 nt running mean at each position on the WT or SS genome was divided by the 15 nt running mean at the corresponding position on the M3 genome. This avoided any instances of division by zero. Only positive-sense reads were used, and library pairs for normalisation were allocated according to replicate number. For the broad-spectrum dataset, guided by the length distribution and phasing plots, reads of lengths 51-52, 54-55 and 57-64 nt were selected for analysis as potential “disome-protected fragments”, and the denominator for normalisation of disome-protected read densities to RPM was calculated using only reads of these lengths. For all plots of read distribution on the viral genome, a +12 nt offset was added to the 5′-end coordinate of the read before plotting, to reflect the inferred position of the ribosomal P site. For disome-protected fragments, this represents the position of the P site of the colliding ribosome. For plots in the main text showing data from only one replicate, replicate 3 was used. Relative adaptiveness values for sense codons to the cellular tRNA pool were downloaded from the Species-Specific tRNA Adaptive Index Compendium (49), for *Cricetulus griseus* as a proxy for *M. auratus*, on 3^rd^ Aug 2020. For the phasing and length distribution quality control plots, only reads which map completely within the CDSs of host mRNAs were included.

Frameshift efficiency (%) is given by the equation 100 × [1-(downstream/upstream)], where downstream and upstream represent the reads per kilobase per million mapped reads (RPKM) values for the respective regions (genomic coordinates: upstream 1368-3943; downstream 4577-7679) after normalisation of WT and SS densities by M3. The percentage of reads in each phase in the upstream, transframe, and downstream regions relative to the 2B* frameshift site were determined using reads with inferred P site positions in the following ranges: upstream 1098-4211, transframe 4247-4273, downstream 4308-7946. Phases were designated relative to the polyprotein reading frame.

## Data Availability

The datasets produced in this study are available in the following databases:

- Structure factors and coordinates (X-ray crystallography): world-wide Protein Data Bank (wwPDB; https://www.ebi.ac.uk/pdbe/; PDB ID **7NBV**)
- Deep sequencing data (Ribosome profiling): ArrayExpress (https://www.ebi.ac.uk/arrayexpress/) accession numbers **E-MTAB-9438** and **E-MTAB-9437**.

## Acknowledgements

We thank the staff of Diamond Light Source beamline I04-1 for assistance with crystal screening and data collection. We thank Janet Deane for assistance with SEC-MALS experiments. We thank Tatyana Koch for expert technical assistance. This research was funded in whole, or in part, by the Wellcome Trust 202797/Z/16/Z. For the purpose of Open Access, the author has applied a CC BY public copyright licence to any Author Accepted Manuscript version arising from this submission. C.H.H. and S.N. are funded by a Wellcome Trust Investigator Award (202797/Z/16/Z) to I.B.. G.M.C. is funded by a Wellcome Trust PhD Studentship (203864/Z/16/Z). S.C.G. is funded by a Sir Henry Dale fellowship (098406/Z/12/B) funded by the Wellcome Trust and the Royal Society. A.E.F. and K.B. are supported by Wellcome Trust (106207) and European Research Council (646891) grants to A.E.F.. N.C. is funded by the Helmholtz Association and European Research Council StG (948636).

## Author contributions

C.H.H. and S.N. cloned, expressed and purified proteins and performed all biochemical experiments. C.H.H. and S.C.G performed crystallography experiments. A.K. and N.C. performed MST experiments. G.C., K.B. and A.E.F performed and analysed ribosomal profiling data. All authors contributed to preparing figures and writing the manuscript.

## Supplementary Figures and Legends

**Figure S1.**
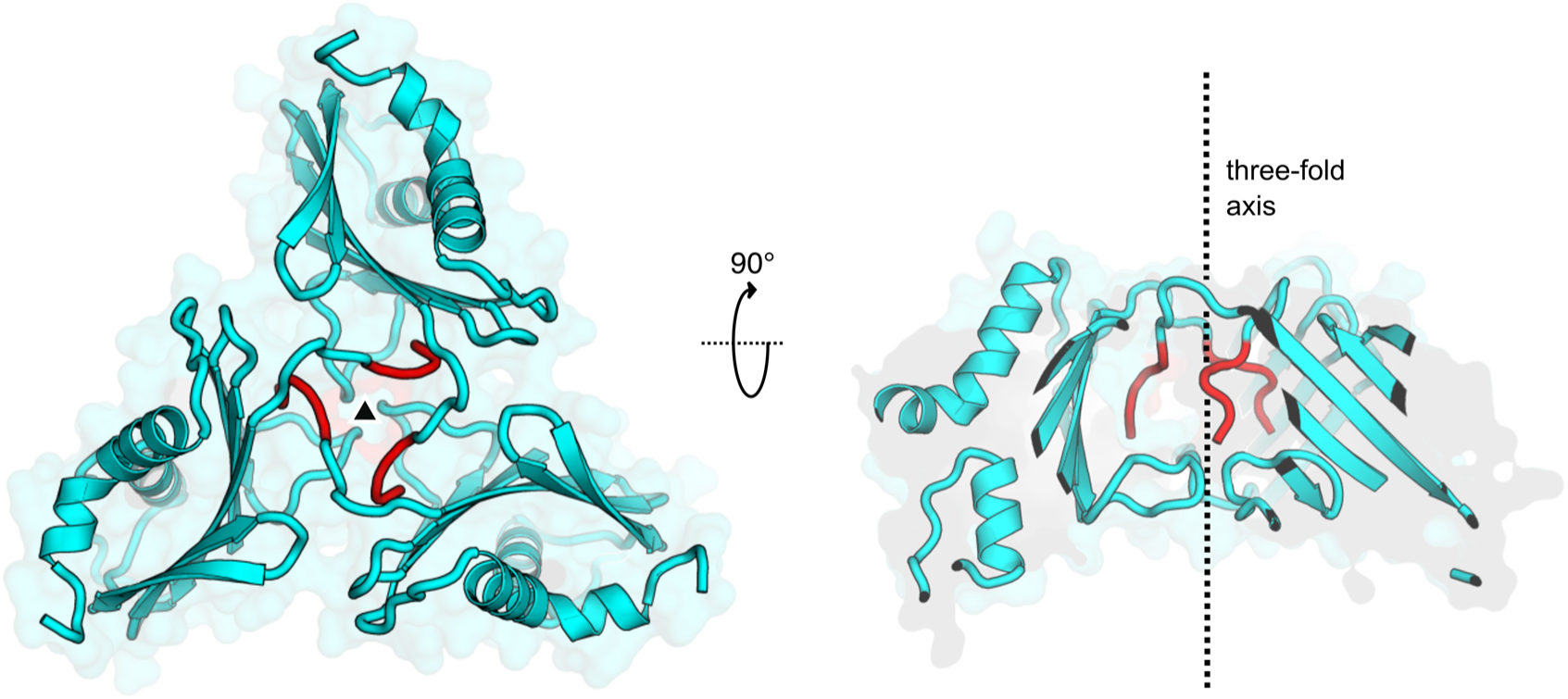
TMEV 2A trimers observed in the crystalline lattice. View of the trimeric protein assembly at the three-fold crystallographic symmetry axis. TMEV 2A molecules are shown as cartoons (cyan) in two orthogonal views, centred around the symmetry axis (black triangle and dotted line). This arrangement may correspond to trimeric assemblies predicted by PDBePISA and observed as a minor species in solution by SEC-MALS (Figure 1B). The interface between subunits is predominantly formed by the N-terminal cloning tag extension (G_-4_P_-3_L_-2_G_-1_S_0_ ; red). The trimeric assembly is therefore not likely to be physiologically relevant.

**Figure S2.**
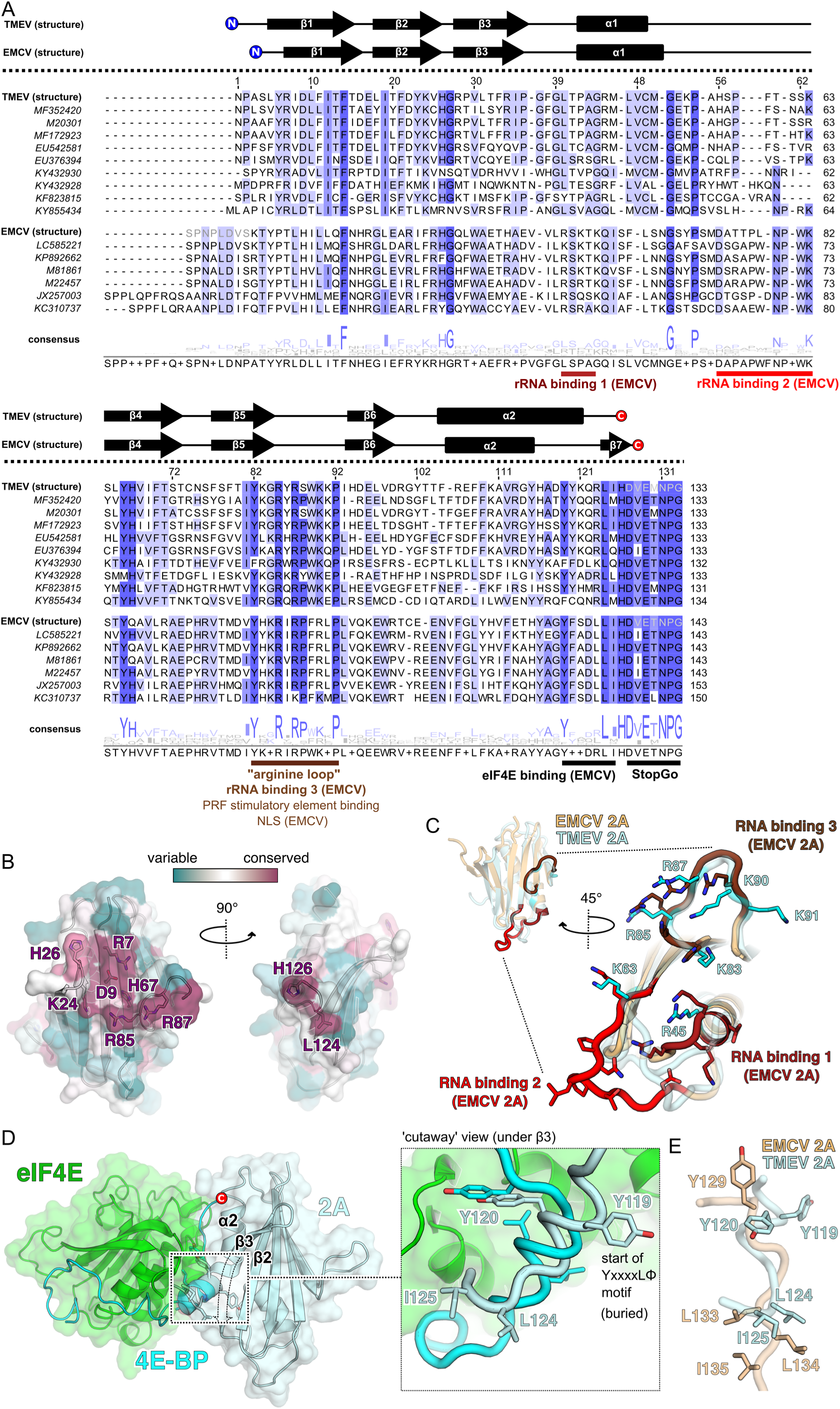
(previous page). Comparison of TMEV 2A structure with orthologue from EMCV reveals a divergent RNA-binding surface and C-terminal region. **A.** Amino acid sequence alignment of selected divergent TMEV-like (upper) and EMCV-like (lower) cardiovirus 2A protein sequences, guided by the structural alignment of EMCV and TMEV 2A proteins. Known secondary structures are indicated above the corresponding sequence for EMCV and TMEV proteins. N-and C-terminal residues not observed in the structures are greyed out. Conservation is highlighted in blue. Local motifs of functional significance are highlighted and annotated. **B.** Surface of TMEV 2A, coloured by conservation from highly conserved (purple) to variable (teal). Highly conserved, surface-accessible residues are shown as sticks. **C.** Comparison of RNA-binding residues in EMCV 2A (28) (beige) with equivalent surface in TMEV 2A (pale blue). Residues involved in RNA binding come from three regions of the EMCV protein as indicated in A (shown as red, crimson and brown sticks). TMEV residues that may be functionally-equivalent are shown as sticks (cyan). **D.** Comparison of the YxxxxLΦ binding motif in 4E-BP1 and putative motif in TMEV 2A. The crystal structure of the complex between eIF4E and 4E-BP1 is shown (green and blue, respectively) with 2A (pale blue) docked via least-squares superposition of the YxxxxLΦ motif. A local surface cutaway, removing a section of β3 (dashed lines), shows the buried location of Y119. *<INSET>* Contrast between the two different helical conformations of the putative 2A YxxxxLΦ motif and the 4E-BP1 YxxxxLΦ motif, in a compact α-helical conformation. **E.** Conformational differences between the C-terminal putative YxxxxLΦ motif in TMEV 2A (pale blue) and EMCV 2A (wheat).

**Figure S3.**
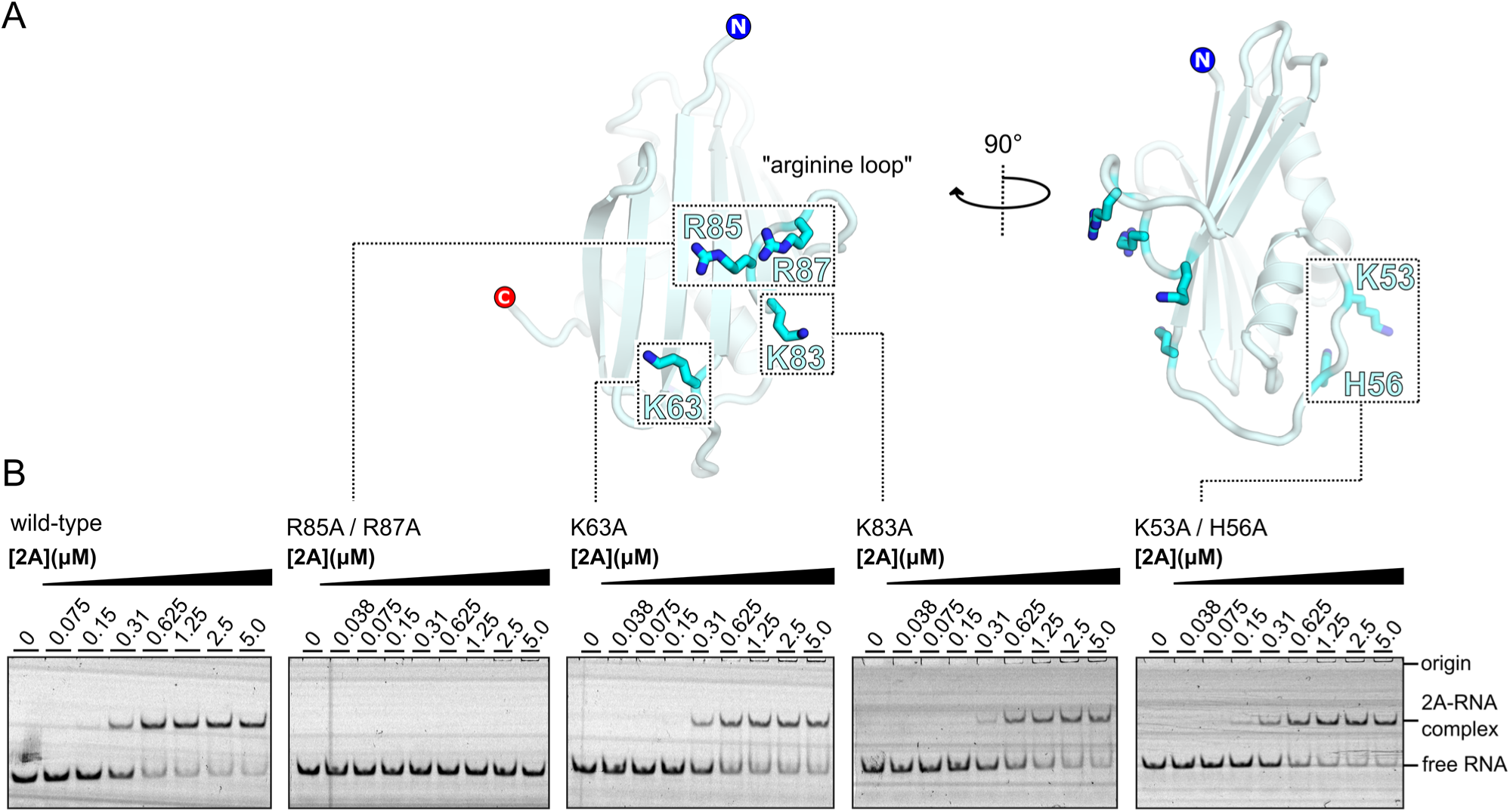
Mutagenesis of putative RNA-binding residues demonstrate the importance of the conserved arginine loop. **A.** Mutagenesis of TMEV residues equivalent to those observed at the RNA binding surface in the EMCV 2A-70S_IC_ structure - see also Figure S2C. The locations of mutations R85A / R87A, K63A, K83A and K53A / H56A are shown as sticks. **B.** EMSA analyses showing effects of the above mutations on stimulatory element RNA binding, compared to a wild-type control. Experiments were conducted with 50 nM Cy5-labelled TMEV 6 RNA and 2A concentrations as indicated between zero and 5.0 μM. Following non-denaturing electrophoresis, fluorescence was imaged using a Typhoon scanner.

**Figure S4.**
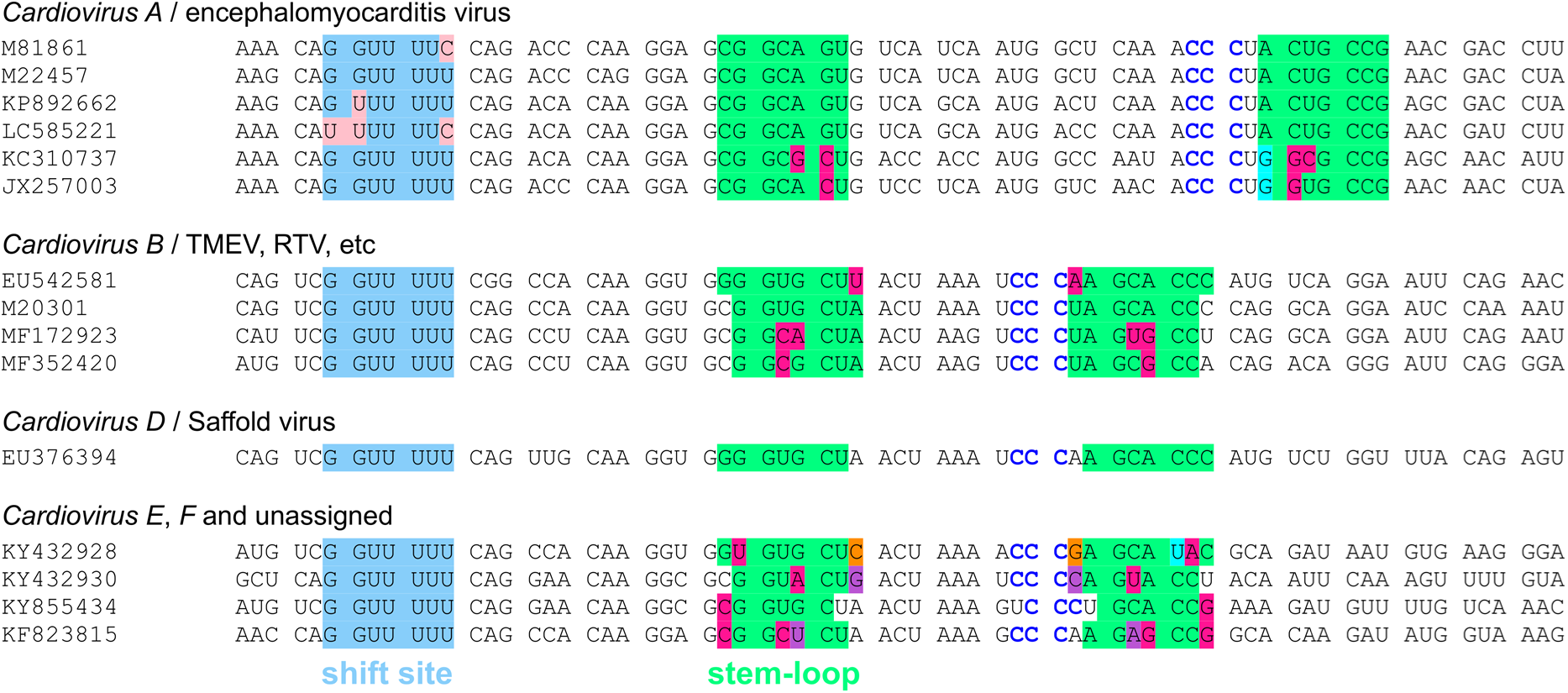
Conservation at the frameshift stimulatory site in cardioviruses. Sequence of the PRF region from representative cardiovirus isolates, showing the highly conserved shift site (light blue) and 3′ RNA stem-loop structure (green). Nucleotide variations in the shift site are indicated in pale pink. Paired substitutions that preserve the predicted structure are highlighted in crimson, purple or orange; single substitutions that – via G:U base-pairing are compatible with the predicted structure – are highlighted in cyan. The conserved CCC loop triplet is shown in blue.

**Figure S5.**
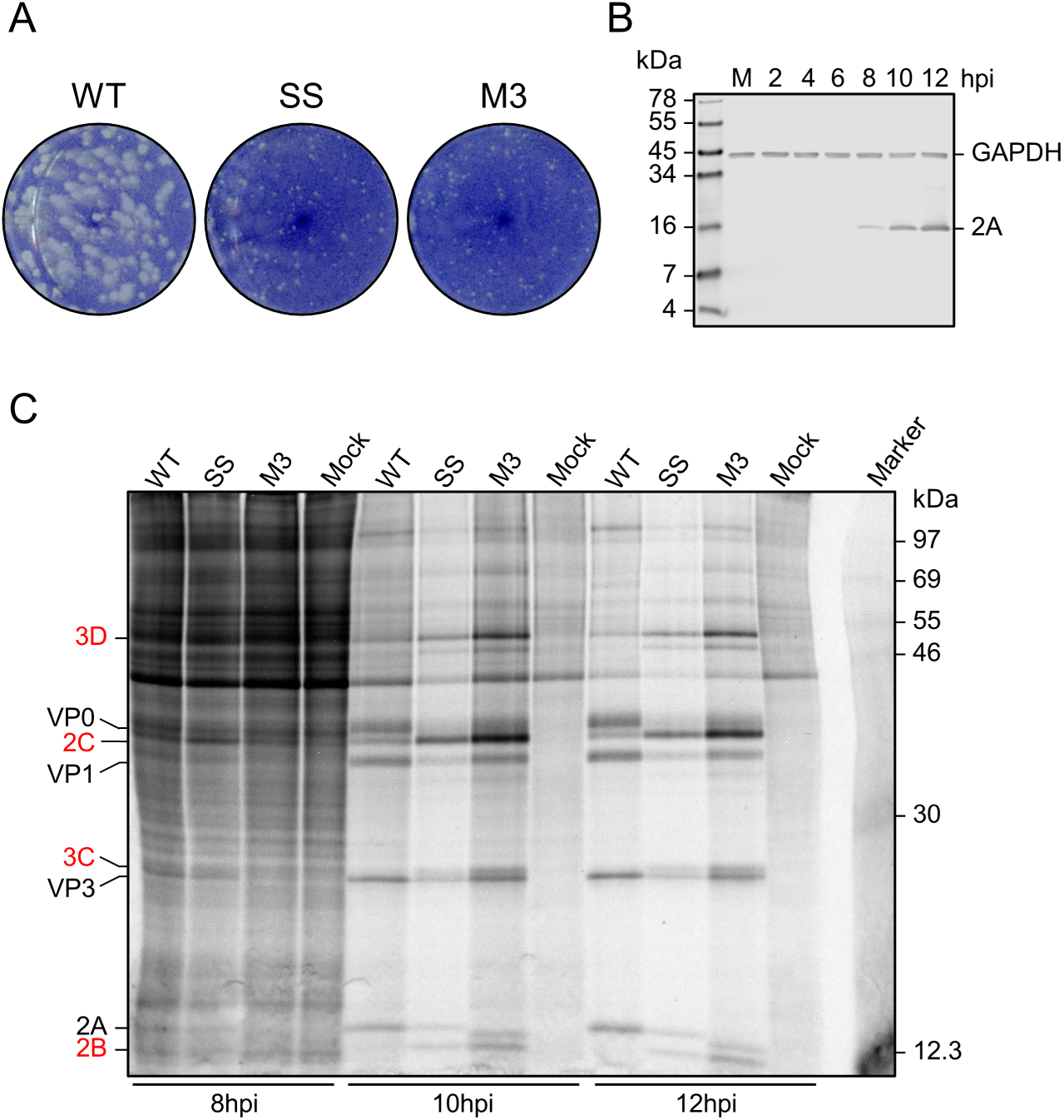
WT and mutant 2A in the context of infection. **A.** Plaque assays of BSR cells infected with WT, SS, or M3 TMEV and fixed at 48 hpi. **B.** Western blot of BSR cells infected with WT TMEV and harvested over a timecourse up to 12 hpi, or mock-infected (M) and harvested immediately. The SS mutant was assayed in parallel and produced similar results (data not shown). **C.** Metabolic labelling of BSR cells infected with WT, SS, or M3 TMEV and harvested over a timecourse of 8-12 hpi. Positions of TMEV proteins are indicated, with those downstream of the frameshift site written in red. At 8 hpi, reliable quantification of viral proteins above the background of host translation was not possible. However, at 10 and 12 hpi, a large proportion of ongoing translation is viral, likely due to virus-induced shut-off of host gene expression, and viral proteins were clearly visible and quantifiable.

**Figure S6.**
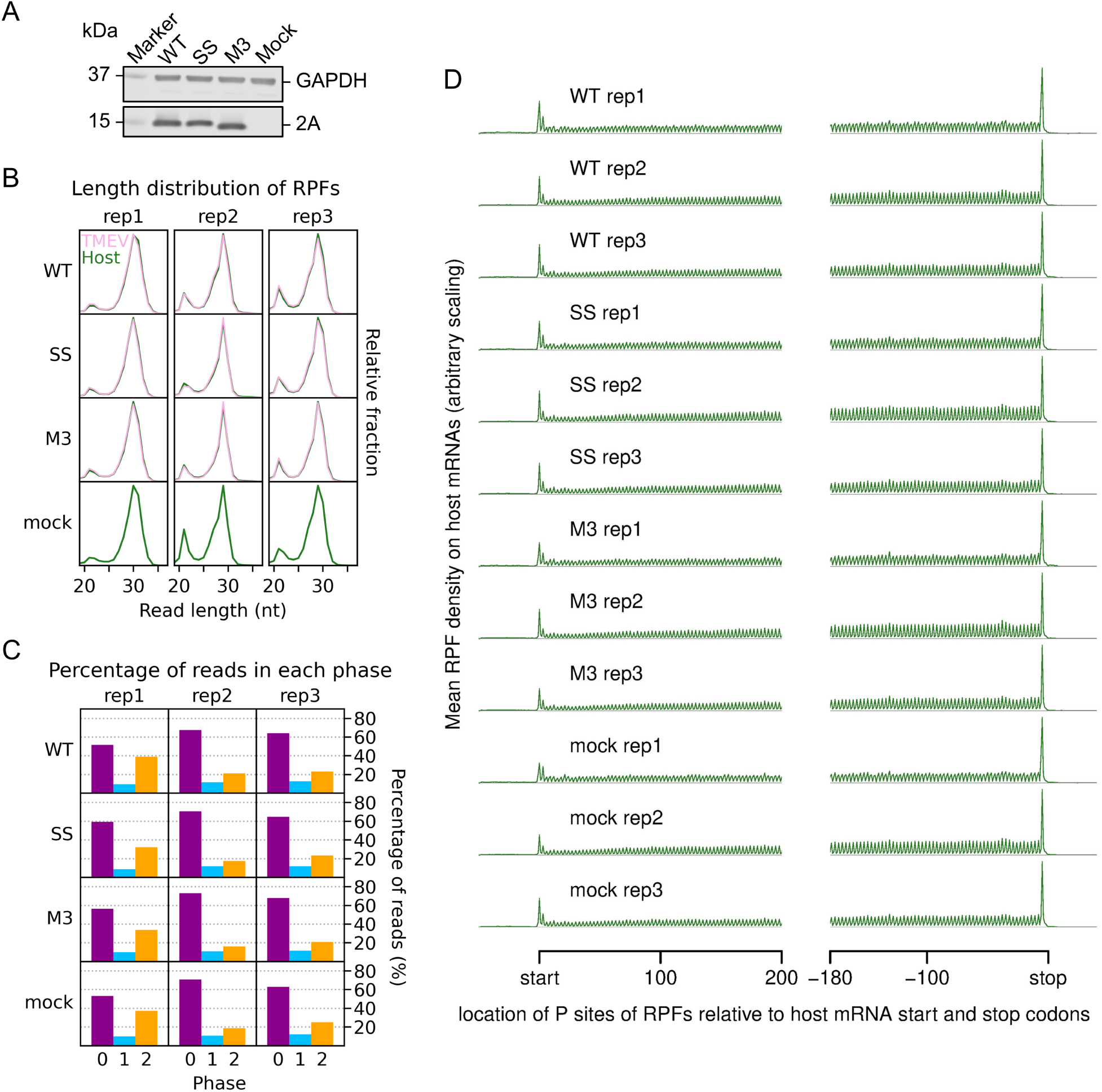
Ribosome profiling quality control analysis indicates high quality data. **A.** Western blot of lysates used for ribosome profiling. Note that the mutant M3 2A migrates slightly faster during gel electrophoresis than the WT protein. **B.** Length distribution of RPFs mapping within host (green) and viral (pink, mock excluded) CDSs in each library. **C.** Percentage of reads (all read lengths) attributed to each phase, from reads mapping within host CDSs. Noting that approximately one third of the reads in replicate 1 libraries were attributed to the −1/+2 phase, we increased the amount of RNase I added to further replicates to improve trimming, yielding the very high proportion of phase 0 reads seen in replicates 2 and 3. Phase compositions of virus CDS-mapping reads closely matched those of host-mapping reads (Figure S8A, upstream and downstream panels). **D.** Distribution of inferred P site positions of host mRNA-mapping reads relative to start and stop codons. Only transcripts with an annotated CDS of at least 150 codons and UTRs of at least 60 nt were included in the analysis, and the total number of reads from all these transcripts mapping to each position was plotted. RPFs map to coding sequences with a triplet periodicity and few RPFs map to the UTRs, particularly the 3′UTR. Typically heightened RPF peaks corresponding to the sites of translation initiation and termination are also observed (32, 87).

**Figure S7.**
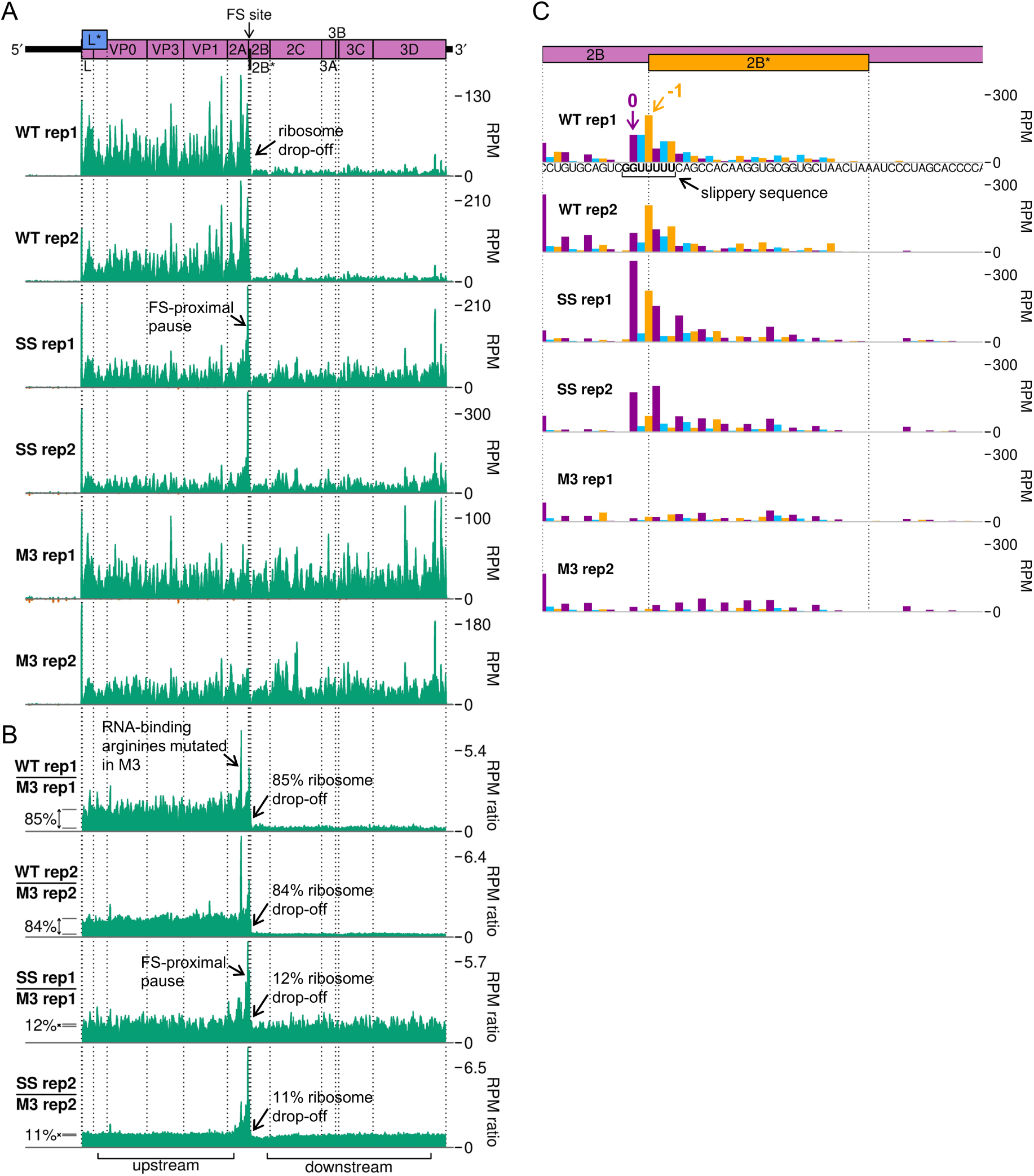
Further replicates of ribosome profiling RPF distribution plots. **A.** RPF distribution on the viral genome, as in main text Figure 6D, for replicates 1 and 2. **B.** Ratio of RPF density on the WT or SS genome normalised by M3, as in main text Figure 6F, for replicates 1 and 2. **C.** RPF distribution at the frameshift site, as in main text Figure 6G, for replicates 1 and 2.

**Figure S8.**
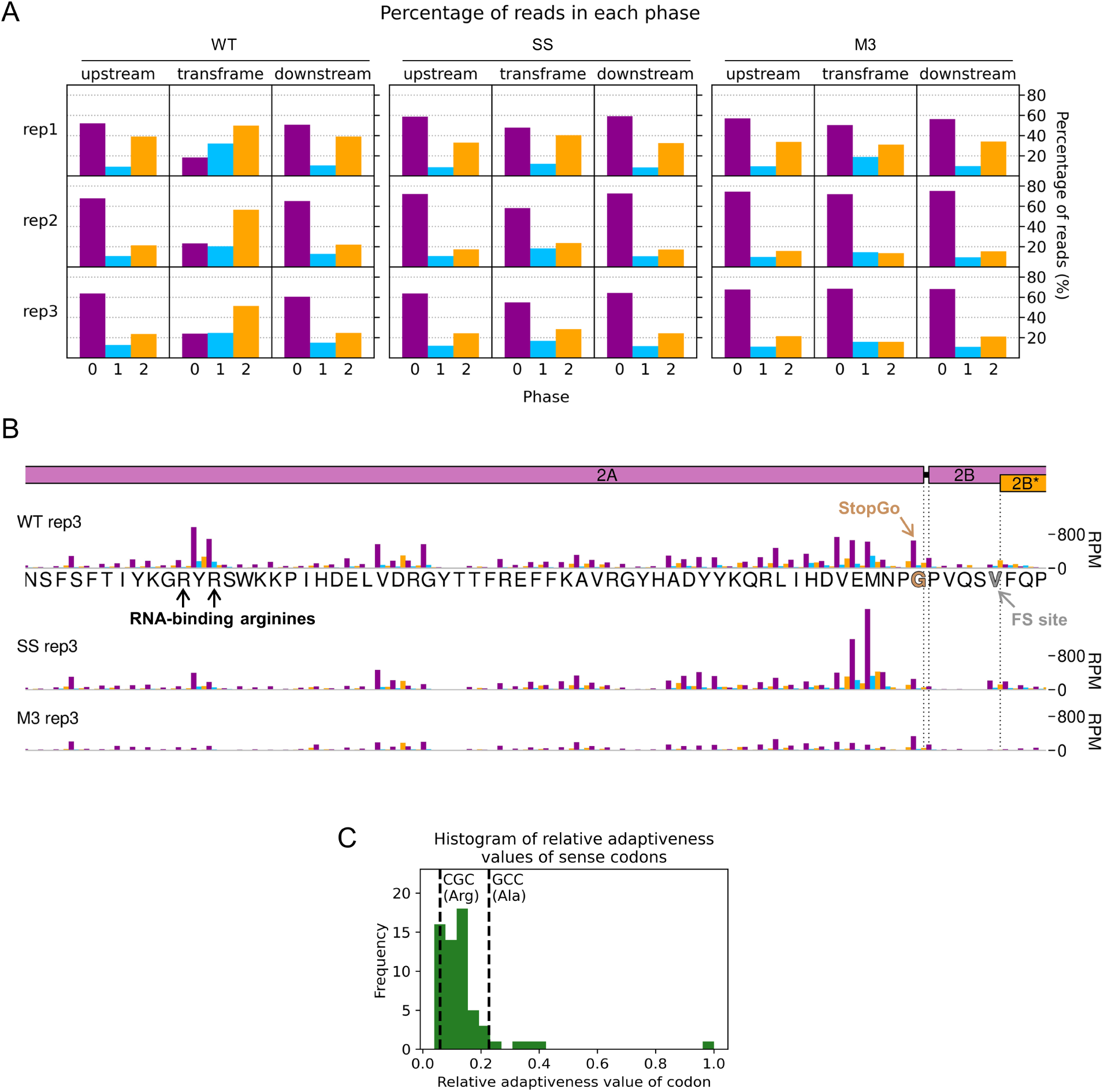
Investigation of the region around the frameshift site by ribosome profiling. **A.** Percentage of reads (all read lengths) attributed to each phase in the regions upstream and downstream of 2B* and in the short 2B/2B* overlap region (“transframe”). **B.** Density of RPFs in the 200 nt up to and including the frameshift site, plotted at inferred P site positions of the ribosomes, and coloured according to RPF phase as in D. The encoded amino acid sequence in this region is underlaid beneath the data in the top panel, and all libraries are set to the same scale on the y axis, with no running mean filter applied. The last amino acid of 2A – after which StopGo-mediated peptide release occurs – is annotated in brown, the P-site of frameshifting (FS) in grey, and the RNA-binding arginines mutated to alanine in M3 in black. **C.** Histogram of relative adaptiveness values of each sense codon to the cellular tRNA pool, defined based on a combination of intracellular tRNA abundance and the strength of the codon-anticodon interaction. Values for the Chinese hamster *Cricetulus griseus* were taken from the Species-Specific tRNA Adaptive Index Compendium (49) as *M. auratus* values were not available. The adaptiveness values of the CGC arginine codons encoding R85 and R87, and the GCC alanine codons to which they are mutated in M3, are indicated by dashed lines.

## Supplementary Tables

**Table S1.** Crystallographic data collection and refinement.

|  | Highest resolution<sup>†</sup> | Structure determination<sup>†</sup> |
| --- | --- | --- |
| **Data collection\*** |  |  |
| Wavelength (Å) | 0.9159 | 0.9159 |
| Space group | *P* 2<sub>1</sub> 3 | *P* 2<sub>1</sub> 3 |
| Cell dimensions |  |  |
| *a* = *b* = *c* (Å) | 73.77 | 74.24 |
| Resolution (Å) | 43.59–1.87 (1.92–1.87) | 52.48–2.11 (2.15–2.11) |
| Unique reflections | 11,355 (577) | 8100 (389) |
| Completeness (%) | 100.0 (100.0) | 100.0 (100.0) |
| Anomalous | - | 100.0 (100.0) |
| Multiplicity | 19.0 (19.8) | 19.4 (20.3) |
| Anomalous |  | 10.3 (10.6) |
| *R*<sub>merge</sub> | 0.113 (1.191) | 0.078 (1.956) |
| *R*<sub>pim</sub> | 0.026 (0.273) | 0.025 (0.629) |
| CC<sub>1/2</sub> | 0.999 (0.644) | 1.00 (0.715) |
| CC<sub>anom</sub> | - | 0.504 (0.002) |
| Mean I/σ(I) | 16.8 (2.8) | 26.9 (1.7) |
| Wilson B (Å<sup>2</sup>) | 26.06 | 46.96 |
| **Refinement\*** |  |  |
| Resolution (Å) | 42.61–1.87 (1.93–1.87) |  |
| Reflections ‡ | 21,277 (2095) |  |
| Working set | 20,162 (1961) |  |
| Test set | 1115 (134) |  |
| *R*<sub>work</sub> | 0.1847 |  |
| *R*<sub>free</sub> | 0.2302 |  |
| No. of atoms |  |  |
| Protein | 1112 |  |
| Solvent | 98 |  |
| Ions | 1 |  |
| Root mean square deviation |  |  |
| Bond lengths (Å) | 0.005 |  |
| Bond angles (°) | 0.890 |  |
| Ramachandran favoured (%) | 100 |  |
| Ramachandran outliers (%) | 0.0 |  |
| Poor rotamers (%) | 0.0 |  |
| Mean B value (Å<sup>2</sup>) | 31.6 |  |

\*Values in parentheses are for the highest resolution shell
† Both crystals were harvested from the same crystallisation drop
‡ I<sup>+</sup> and I<sup>-</sup> were treated as separate reflections in phenix.refine

**Table S2.** Number of ribosome profiling reads assigned to each category. The “_broad” samples are the broad-spectrum libraries, for which 35-65 nt fragments were purified. Reads under 19 nt long (for monosome libraries) or under 35 nt long (for broad-spectrum libraries) were defined as too short. Some categories are not applicable to the broad-spectrum libraries due to minor differences in computational processing (detailed in Methods).

| **Virus** | **Replicate** | **Total reads** | **Too short** | **Adapter-only** | **No adapter** | **PCR duplicates** | **rRNA** |
| --- | --- | --- | --- | --- | --- | --- | --- |
| WT | 1 | 1790088 | 61648 | 876 | 12076 | 28721 | 586963 |
| WT | 2 | 26408581 | 2340239 | 297018 | 116064 | 1834921 | 2926112 |
| WT | 3 | 28688961 | 2756571 | 280096 | 406493 | 2627172 | 4719318 |
| SS | 1 | 2392994 | 34429 | 1859 | 9565 | 61705 | 505571 |
| SS | 2 | 25027728 | 2854707 | 353732 | 126430 | 1456929 | 2342432 |
| SS | 3 | 21788351 | 1927596 | 320336 | 503529 | 2258152 | 3634096 |
| M3 | 1 | 2290607 | 149115 | 6052 | 18977 | 61127 | 619963 |
| M3 | 2 | 27308973 | 1702862 | 157048 | 138108 | 1654696 | 2986823 |
| M3 | 3 | 30504465 | 2009396 | 515817 | 438248 | 3248459 | 5122599 |
| Mock | 1 | 1914259 | 171017 | 1448 | 27268 | 34259 | 495928 |
| Mock | 2 | 26398293 | 2345528 | 117083 | 120712 | 1286218 | 2852963 |
| Mock | 3 | 27548554 | 1532098 | 145612 | 608503 | 2112106 | 4820304 |
| WT_broad | 3 | 10624342 | 2528401 | 1155 |  |  | 2546503 |
| M3_broad | 3 | 7230428 | 1686167 | 1182 |  |  | 2090553 |
| mock_broad | 3 | 9730736 | 1814114 | 950 |  |  | 3247839 |

| **Virus** | **Replicate** | **vRNA** | **mRNA** | **ncRNA** | **mtDNA** | **gDNA** | **BL21 E. coli** |
| --- | --- | --- | --- | --- | --- | --- | --- |
| WT | 1 | 60322 | 371302 | 71085 | 1909 | 81527 | 2196 |
| WT | 2 | 789308 | 3476556 | 1470387 | 119371 | 930513 | 8738367 |
| WT | 3 | 1299982 | 5671747 | 2271869 | 61429 | 1890651 | 914704 |
| SS | 1 | 198472 | 700308 | 189411 | 7642 | 139138 | 6991 |
| SS | 2 | 769385 | 2293504 | 980217 | 49001 | 527344 | 11085126 |
| SS | 3 | 1136138 | 4143979 | 1661076 | 64024 | 1486102 | 650083 |
| M3 | 1 | 102620 | 485732 | 111611 | 2746 | 129914 | 5070 |
| M3 | 2 | 946929 | 3136082 | 1210389 | 56677 | 837885 | 12118144 |
| M3 | 3 | 1669867 | 6825341 | 2494147 | 105091 | 2404980 | 586000 |
| Mock | 1 | 161 | 481985 | 88888 | 3701 | 105409 | 4205 |
| Mock | 2 | 426 | 4817616 | 904360 | 26897 | 1216025 | 10404660 |
| Mock | 3 | 742 | 9839798 | 1651028 | 67215 | 1765457 | 427314 |
| WT_broad | 3 | 97581 | 534229 | 1322626 | 271988 | 413111 | 485534 |
| M3_broad | 3 | 20310 | 327007 | 783510 | 268166 | 270980 | 110028 |
| mock_broad | 3 | 31 | 349897 | 1038588 | 294835 | 295848 | 163860 |

**Table S3.** Phosphorimager signals for viral-specific bands on metabolic labelling gels.

**10 hpi replicate 1**
| **Protein** | **WT** | **SS** | **M3** | **Carried forward for calculations?** |
| --- | --- | --- | --- | --- |
| 3D | 2395.08 | 2470.9 | 6697.62 | Yes |
| VP0 | 3099.31 |  |  | No - band too faint for reliable quantification in mutant lanes |
| 2C | 2741.89 | 5435.87 | 10800.7 | Yes |
| VP1 | 2976.72 | 1511.32 | 3126.03 | Yes |
| 3C |  | 861.87 | 1954.94 | No - band too faint for reliable quantification in mutant lanes |
| VP3 | 2703.24 | 1402.52 | 3178.16 | Yes |
| 2A | 2325.6 | 1083.08 | 1736.97 | No - methionine-normalised results suggest difference in turnover rate between WT and mutant 2A |
| 2B | 853.34 | 1141.5 | 2323.76 | Yes |

**10 hpi replicate 2**
| **Protein** | **WT** | **SS** | **M3** | **Carried forward for calculations?** |
| --- | --- | --- | --- | --- |
| 3D | 28715.83 | 26525.91 | 67503.74 | Yes |
| VP0 | 32548 |  |  | No - band too faint for reliable quantification in mutant lanes |
| 2C | 25909.88 | 39483.35 | 90913.71 | Yes |
| VP1 | 28175.9 | 13058.42 | 31910.11 | Yes |
| 3C |  | 9944.51 | 20698.75 | No - band too faint for reliable quantification in WT lane |
| VP3 | 23789.64 | 12568.31 | 28983.54 | Yes |
| 2A | 15459.04 | 8312.66 | 13068.13 | No - methionine-normalised results suggest difference in turnover rate between WT and mutant 2A |
| 2B |  | 9735.73 | 16639.4 | No - band too faint for reliable quantification in WT lane |

**10 hpi replicate 3**
| **Protein** | **WT** | **SS** | **M3** | **Carried forward for calculations?** |
| --- | --- | --- | --- | --- |
| 3D | 1272.31 | 1287.21 | 2652.87 | Yes |
| VP0 | 1577.01 |  |  | No - band too faint for reliable quantification in mutant lanes |
| 2C | 1116.87 | 2546.23 | 3469.65 | Yes |
| VP1 | 1155.83 | 729.01 | 889.83 | Yes |
| 3C |  |  | 600.71 | No - band too faint for reliable quantification in WT lane |
| VP3 | 1089.86 | 608.08 | 1074.88 | Yes |
| 2A |  |  |  | No - band not resolvable from dye front |
| 2B |  |  |  | No - band not resolvable from dye front |

**12 hpi replicate 1**
| Protein | WT | SS | M3 | Carried forward for calculations? |
| --- | --- | --- | --- | --- |
| 3D | 14521.16 | 21433.11 | 43338.38 | Yes |
| VP0 | 31017.26 |  |  | No - band too faint for reliable quantification in mutant lanes |
| 2C | 19216.2 | 33218.68 | 63307.42 | Yes |
| VP1 | 28838.43 | 11491.95 | 20683.21 | Yes |
| 3C |  | 9132.88 | 16079.93 | No - band too faint for reliable quantification in WT lane |
| VP3 | 23912.68 | 11430.74 | 20631.82 | Yes |
| 2A | 15548.96 | 7338.38 | 8650.82 | No - methionine-normalised results suggest difference in turnover rate between WT and mutant 2A |
| 2B | 5747.61 | 7299.89 | 11206.57 | Yes |

**12 hpi replicate 2**
| Protein | WT | SS | M3 | Carried forward for calculations? |
| --- | --- | --- | --- | --- |
| 3D | 945.31 | 2963.3 | 7539.19 | Yes |
| VP0 | 2397.33 |  |  | No - band too faint for reliable quantification in mutant lanes |
| 2C | 1470.08 | 5552.5 | 10498.07 | Yes |
| VP1 | 2248.87 | 1656.4 | 3280.92 | Yes |
| 3C |  | 963.09 | 1771.51 | No - band too faint for reliable quantification in WT lane |
| VP3 | 2434.33 | 1869.35 | 3682.06 | Yes |
| 2A | 1847.17 | 1285.68 | 1844.53 | No - methionine-normalised results suggest difference in turnover rate between WT and mutant 2A |
| 2B | 476.88 | 1350.26 | 2322.62 | Yes |

**12 hpi replicate 3**
| Protein | WT | SS | M3 | Carried forward for calculations? |
| --- | --- | --- | --- | --- |
| 3D | 664.88 | 7645.55 | 8779.04 | Yes |
| VP0 | 1760.34 |  |  | No - band too faint for reliable quantification in mutant lanes |
| 2C | 934.12 | 7949.59 | 10576.14 | Yes |
| VP1 | 1515.47 | 2134.25 | 2391.4 | Yes |
| 3C |  | 952.4 | 621.45 | No - band too faint for reliable quantification in WT lane |
| VP3 | 1619.9 | 2328.56 | 2969.14 | Yes |
| 2A |  |  |  | No - band not resolvable from dye front |
| 2B |  |  |  | No - band not resolvable from dye front |

